# TET loss of function selects for aneuploidies that confer a proliferative advantage

**DOI:** 10.64898/2026.09.27.754858

**Authors:** Isaac F. López-Moyado, Kazumasa Suzuki, Lot Hernández-Espinosa, Xiang Li, Daniëlle Koot, René Wardenaar, Floris Foijer, Anjana Rao

## Abstract

Mutations in the Ten-Eleven Translocation (TET) family of DNA dioxygenases are among the most common epigenetic alterations in clonal hematopoiesis and hematologic malignancies, yet how TET loss of function reshapes genome evolution during cellular expansion remains to be fully understood. Here we show that loss of *Tet2/3* in invariant natural killer T (iNKT) cells drives aggressive lymphoid expansion that is consistently associated with recurrent chromosome 17 trisomy, a dominant aneuploid state observed across more than ten independently derived tumors following transplantation into immunocompetent recipient mice. Single-cell whole-genome sequencing of TET-deficient iNKT cells induced to proliferate in culture with antigen and IL-2 stimulation revealed that rare pre-existing cells with chromosome 17 trisomy appeared soon after *Tet* gene deletion in the thymus, together with cells harboring other copy number alterations, but only the cells with chromosome 17 trisomy were progressively enriched during further expansion. Similarly, inducibly *Tet*-deficient mouse embryonic stem cells (mESCs) showed selection for aneuploid states that were present prior to *Tet* gene deletion, including trisomy of chromosome 6. Reanalysis of independent datasets from *Dnmt3a*-deficient mouse lymphomas identified recurrent chromosome gains, including chromosome 17 trisomy. Our results establish *TET* and *DNMT3A* deficiency as drivers of karyotypic evolution, and suggest that disruption of DNA methylation/ demethylation pathways through either *TET* or *DNMT3A* deletion promotes chromosomal instability, in part by facilitating the emergence and selection of cryptic aneuploid subclones during clonal expansion.

## Introduction

DNA methylation is a fundamental epigenetic modification required for the regulation of gene expression, chromatin structure, and cellular identity (Smith et al., 2025). The balance between DNA methylation and demethylation is maintained by the opposing activities of DNA methyltransferases and ten-eleven translocation (TET) dioxygenases. The TET family proteins TET1, TET2, and TET3 catalyze the iterative oxidation of 5-methylcytosine and play critical roles in active and passive DNA demethylation, developmental gene regulation, and maintenance of chromatin organization (Ko et al., 2010; López-Moyado et al., 2024; Tahiliani et al., 2009; Wu & Zhang, 2017). Disruption of these pathways is strongly associated with human disease. In particular, loss-of-function mutations in TET2 are among the most common genetic alterations detected in clonal hematopoiesis, myeloid malignancies, and lymphoid cancers (Delhommeau et al., 2009; Langemeijer et al., 2009; Lemonnier et al., 2012; Lio et al., 2019; Odejide et al., 2014), whereas mutations in DNMT3A, a de novo DNA methyltransferase that functionally opposes TET-mediated demethylation, are similarly associated with age-related clonal expansion and hematologic transformation (Brunetti et al., 2017; Couronné et al., 2012; Jaiswal et al., 2014; Xie et al., 2014). These observations highlight the role of DNA methylation—and more generally, cytosine modifications—as a central regulator of hematopoietic cell fate and malignant evolution.

Previous studies have demonstrated that TET proteins are essential for normal lymphocyte development and immune homeostasis (López-Moyado et al., 2024). We previously showed that combined deletion of *Tet2* and *Tet3* in T cells (using a CD4-Cre recombinase approach) results in rapid clonal expansion of invariant natural killer T (iNKT) cells (López-Moyado et al., 2019; Tsagaratou et al., 2017), a specialized T-cell lineage that expresses an invariant T cell receptor alpha (TCRα) chain, recognizes lipid antigens presented by the non-classical MHC molecule CD1d, and are not deleted in the thymus (Bendelac et al., 2007; Engel & Kronenberg, 2012). *Tet2/3*-deficient mice develop an aggressive and fully penetrant iNKT cell lymphoma that can be serially transplanted into recipient animals, including immunocompetent hosts (Tsagaratou et al., 2017). Disease propagation depends on CD1d expression in recipient mice, indicating that antigen-driven proliferation contributes to malignant expansion (Tsagaratou et al., 2017). These studies established a critical role for TET proteins in restraining aberrant lymphocyte proliferation, but the genomic alterations that accompany disease progression remain incompletely understood.

A growing body of evidence suggests that in addition to their well-established roles in transcriptional regulation, TET proteins may influence genome stability (Georges et al., 2022; Gioulbasani et al., 2024). Aneuploidy and copy-number alterations are hallmarks of cancer and are increasingly recognized as adaptive features of clonal evolution rather than merely passive consequences of genomic instability (Baker et al., 2024; Sdeor et al., 2024; Shoshani et al., 2021; Trakala et al., 2021). Recurrent chromosome gains frequently emerge during tumor progression, where selective pressures can favor karyotypic variants that confer proliferative or survival advantages (Baker et al., 2024; Sdeor et al., 2024). Consistent with this concept, we recently demonstrated that acute deletion of all three *Tet* genes in mouse embryonic stem cells leads to chromosome segregation defects, micronucleus formation, chromatin bridges, lagging chromosomes, and increased frequencies of aneuploidy, a phenotype attributed at least in part to transcriptional downregulation of *Khdc3* associated with promoter hypermethylation (Georges et al., 2022). These observations raised the possibility that disruption of DNA methylation homeostasis may promote not only transcriptional dysregulation but also the generation of heritable chromosomal alterations capable of influencing cellular fitness.

The mechanisms linking epigenetic dysregulation to chromosomal evolution remain poorly understood. DNA hypomethylation has long been associated with genome instability, chromosome segregation defects, and recurrent aneuploidy in both experimental models and human cancers (Eden et al., 2003; Gaudet et al., 2003; Salinas-Luypaert et al., 2025). Loss of methylation within repetitive and heterochromatic centromeric regions can impair chromosome organization and centromere function, increasing the probability of chromosome mis-segregation during cell division (Salinas-Luypaert et al., 2025). Importantly, mutations affecting both TET proteins and DNA methyltransferases are recurrent drivers of clonal hematopoiesis (Jaiswal & Ebert, 2019), suggesting that dysregulation of DNA methylation may create permissive conditions for the emergence and selection of advantageous chromosomal states. Whether recurrent aneuploidies arise during expansion of *Tet*- deficient lymphoid populations, and whether similar chromosomal alterations emerge following perturbation of other epigenetic regulators such as *de novo* DNA methyltransferases, remains unknown.

Here, we investigated the genomic evolution of *Tet2/3*-deficient iNKT cells during transplantation and *in vitro* expansion using bulk and single-cell genomic approaches. We identify recurrent trisomy of mouse chromosome 17 as a predominant chromosomal alteration acquired during expansion of *Tet2/3*-deficient iNKT cells upon transplantation, and show that cells carrying chromosome 17 trisomy exist as rare subclones that become selectively enriched during proliferation. Extending these findings beyond the T cell lineage, we demonstrate that *Tet*-deficient embryonic stem cells similarly undergo selection for recurrent aneuploid subclones, albeit involving a distinct chromosome—chromosome 6. Finally, reanalysis of independent *Dnmt3a*-deficient mouse lymphoma datasets also revealed recurrent chromosome gains, including chromosome 17 trisomy, indicating that perturbations of either the DNA methylation or demethylation machinery can converge to produce related patterns of karyotypic evolution. Together, our findings support a model in which disruption of DNA methylation control promotes the emergence and selective expansion of cryptic aneuploid subclones during cellular proliferation.

## Results

### *Tet2/3*-deficient iNKT cells acquire recurrent chromosome 17 trisomy following transplantation

We previously demonstrated that combined deletion of *Tet2* and *Tet3* in T cells (*Tet2^-/-^ Tet3^fl/fl^ CD4-Cre*; *Tet2/3 DKO*) results in rapid expansion of invariant natural killer T (iNKT) cells and the development of an aggressive transplantable iNKT cell malignancy (Tsagaratou et al., 2017). *Tet2/3 DKO* mice exhibit marked accumulation of iNKT cells beginning early after birth and all mice succumb to lymphoma within 8 weeks (Tsagaratou et al., 2017). Moreover, purified *Tet2/3*-deficient iNKT cells efficiently transfer disease to secondary fully immunocompetent recipients, and disease propagation depends on CD1d expression in the recipient mice, indicating that antigen/MHC-driven stimulation is a major driver of malignant expansion (Tsagaratou et al., 2017).

To investigate whether malignant progression of *Tet2/3*-deficient iNKT cells is associated with recurrent large-scale genomic alterations, we performed genome-wide analyses of donor-derived *Tet2/3 DKO* iNKT cells following transplantation and expansion in recipient mice (**Fig. 1a**) (López-Moyado et al., 2019). A total of ten independent tumors were analyzed using multiple orthogonal genomic approaches, yielding whole-genome bisulfite sequencing (WGBS; n = 2), Hi-C (n = 3) and whole-genome sequencing (WGS; n = 5) datasets (López-Moyado et al., 2019; Tsagaratou et al., 2017). Despite the distinct experimental methodologies, copy-number profiles derived from all three platforms revealed extensive aneuploidy, including recurrent whole-chromosome gains together with less frequent copy number alterations. Notably, chromosome 17 trisomy was detected in all ten expanded *Tet2/3 DKO* iNKT-cell tumors analyzed, making it the only whole-chromosome alteration shared across all samples (**Fig. 1b**).

**Figure 1.**
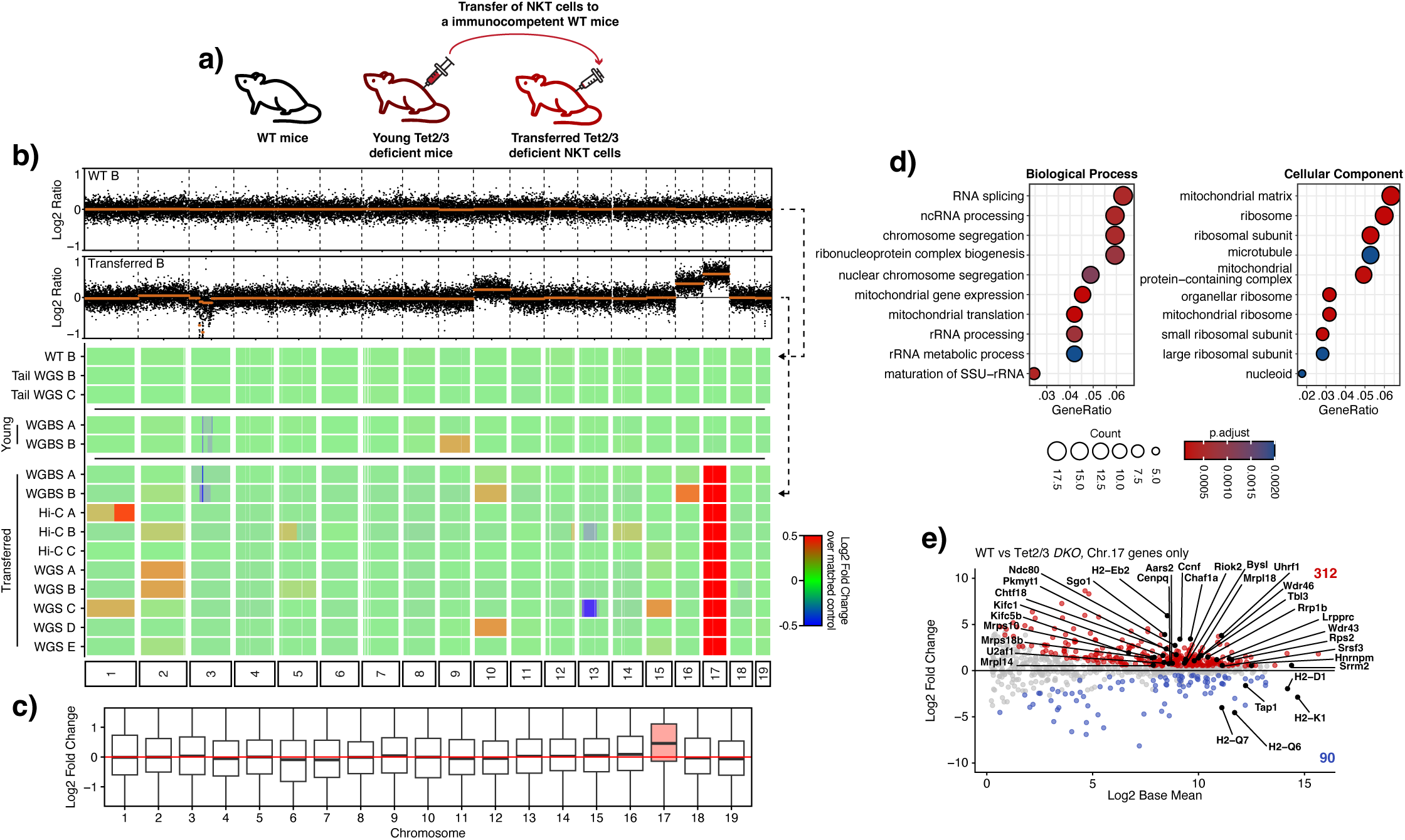
Recurrent chromosome 17 trisomy in *Tet2/3*-deficient iNKT-cell lymphoma. (a) Experimental scheme. Purified *Tet2/3 DKO* iNKT cells isolated from donor mice were transferred into immunocompetent recipient animals and analyzed 9-12 wk following malignant expansion. (b) Genome-wide copy-number profiles of wild-type (WT) iNKT cells, *Tet2/3 DKO* iNKT cells prior to transplantation (*Young*), and *Tet2/3 DKO* iNKT cells recovered after transplantation (*Transferred*). Data were derived from whole-genome sequencing (WGS), whole-genome bisulfite sequencing (WGBS), and Hi-C datasets (and previously published in (López-Moyado et al., 2019; Tsagaratou et al., 2017)). Each point represents a 100-kb genomic window. Red indicates copy-number gain and blue indicates copy-number loss. Recurrent gain of chromosome 17 is observed in independently derived transplanted tumors. (c) Chromosome-level distribution of gene-expression changes in transplanted *Tet2/3 DKO* iNKT cells relative to WT splenic iNKT cells. Boxplots show log_2_ fold-change values for all expressed genes on each chromosome. Chromosome 17 genes display a global increase in expression consistent with chromosome gain. (d) Gene Ontology enrichment analysis of chromosome 17 genes significantly upregulated (log_2_FC>0.5 and *P*-adjusted value <0.05 using the Benjamini–Hochberg test) in transplanted *Tet2/3 DKO* iNKT cells. Enriched categories include RNA splicing, rRNA processing, chromosome segregation, and mitochondrial gene expression. (e) MA plot of RNA-seq data showing differential gene expression of chromosome 17 genes in *Tet2/3 DKO* iNKT cells recovered after transplantation relative to WT iNKT cells. Genes meeting the criteria for differential expression (|log_2_FC|>0.5 and *P*-adjusted value <0.05 using the Benjamini–Hochberg test) are indicated in red (upregulated) or blue (downregulated).

In addition to chromosome 17 gain, many tumors exhibited partial and often recurrent trisomies affecting other chromosomes (e.g., chr1, chr2, chr10), as well as focal gains and losses at multiple genomic loci (**Fig. 1b**). A related study of expanded *TET*-deficient T cells using the same mouse strain (Gioulbasani et al., 2024) also reported recurrent chromosome 17 amplification following one or two rounds of transfer into recipient mice, together with recurrent aneuploidies involving chromosomes 2, 15 and 16 after the first transfer and chromosomes 6 and 15 after the second transfer (Gioulbasani et al., 2024). However, no other chromosomal alteration in this study displayed the same degree of consistency across independent tumors as the chromosome 17 trisomy observed in our cohort. Together, these findings identify chromosome 17 gain as a recurrently selected event associated with malignant expansion of *Tet2/3*-deficient iNKT cells *in vivo*.

To determine whether chromosome 17 gain was already present before transplantation, we analyzed WGBS data from bulk populations of iNKT cells isolated from two young *Tet2/3 DKO* donor mice prior to transfer. Although one donor sample displayed a partial trisomy of chromosome 9, neither sample exhibited evidence of chromosome 17 gain (**Fig. 1b**). Thus, chromosome 17 trisomy is not a prominent feature present in a majority of pre-transfer *Tet2/3 DKO* iNKT cell populations. These observations suggest that chromosome 17 trisomy either arises during tumor progression or is present at frequencies below the detection threshold of bulk analyses in pre-transfer donor populations. In either case, cells with chromosome 17 trisomy must subsequently undergo strong positive selection following transplantation into immunocompetent recipient mice.

Consistent with chromosome 17 trisomy, genes located on chromosome 17 showed a chromosome-wide increase in expression relative to genes located on other chromosomes (**Fig. 1c**). Gene Ontology analysis of chromosome 17 genes significantly upregulated in transferred and expanded *Tet2/3 DKO* iNKT cells revealed enrichment for pathways involved in rRNA processing, RNA splicing, chromosome segregation, and mitochondrial gene expression (**Fig. 1d,e**; **Supplementary Tables 1 and 2**). These categories included genes encoding proteins involved in ribosome biogenesis and rRNA processing (*Wdr43*, *Wdr46*, *Rrp1b*, *Tbl3*, *Riok2*, *Rps2*, *Bysl*), RNA splicing (*Srsf3*, *U2af1*, *Hnrnpm*, *Srrm2*), chromosome segregation regulators (*Ndc80*, *Sgo1*, *Cenpq*, *Kifc1*, *Kifc5b*, *Ccnf*, *Chtf18*, *Pkmyt1*), and components of the mitochondrial translation machinery (*Mrps18b*, *Mrps10*, *Mrpl18*, *Mrpl14*, *Lrpprc*). The preferential enrichment for these proliferative and biosynthetic programs suggests that iNKT cell expansion in *Tet2/3 DKO* mice may depend on increased dosage, and hence increased expression, of a subset of chromosome 17 genes that support the growth and expansion of malignant cells.

Increased chromosome 17 copy number did not result in increased expression of all chromosome 17 genes. Genes involved in antigen processing and presentation, including major histocompatibility complex (MHC) genes located on chromosome 17, were significantly downregulated in transplanted *Tet2/3 DKO* iNKT cells compared to wild type (WT) controls (**Fig. 1e**; **Suppl. Fig. S1**). Of note, the gene encoding CD1d, required for iNKT cell antigen recognition, is located on mouse chromosome 3. Thus, despite the increased copy number of chromosome 17, MHC gene expression was generally reduced (**Suppl. Fig. S1b**; **Supplementary Table 3**), indicating that transcriptional regulation can override the dosage effect associated with chromosome gain. This observation further suggests that the biological consequences of chromosome 17 gain are unlikely to result simply from a global increase in chromosome-wide gene dosage, but rather may depend on specific genes whose expression is preferentially increased. We are currently investigating which chromosome 17 genes, individually or in combination, mediate the observed proliferative advantage.

Together, these findings identify chromosome 17 trisomy as a highly recurrent genomic feature of *Tet2/3*- deficient iNKT-cell lymphoma and suggest that increased dosage of chromosome 17 genes contributes to the selective expansion of malignant *TET*-deficient clones.

### Expansion of *Tet2/3*-deficient iNKT cells can be recapitulated *in vitro*

The recurrent emergence of chromosome 17 trisomy and other chromosomal copy number alterations (CNAs) following transplantation suggested that these genomic alterations conferred a selective advantage to *Tet2/3*- deficient iNKT cells during *in vivo* expansion in recipient mice. To determine whether this enhanced proliferative capacity could be recapitulated under defined conditions in the absence of an *in vivo* microenvironment or stimulus, we established an *in vitro* culture system for primary iNKT cells. Splenocytes isolated from *Tet2^fl/fl^ Tet3^fl/fl^ CD4-Cre* (*Tet2/3 DKO*) or *Tet2^fl/fl^ Tet3^fl/fl^*(*Tet2/3 floxed*, WT) mice were cultured in the presence of IL-2 and an α-Galactosylceramide (α-GalCer)-loaded CD1d tetramer for six days, followed by continued culture in IL- 2 alone (**Fig. 2a**). α-GalCer is a glycoplipid antigen that potently activates iNKT cells (Kawano et al., 1997). The frequency of iNKT cells was monitored by flow cytometry at culture initiation and throughout the expansion period.

**Figure 2.**
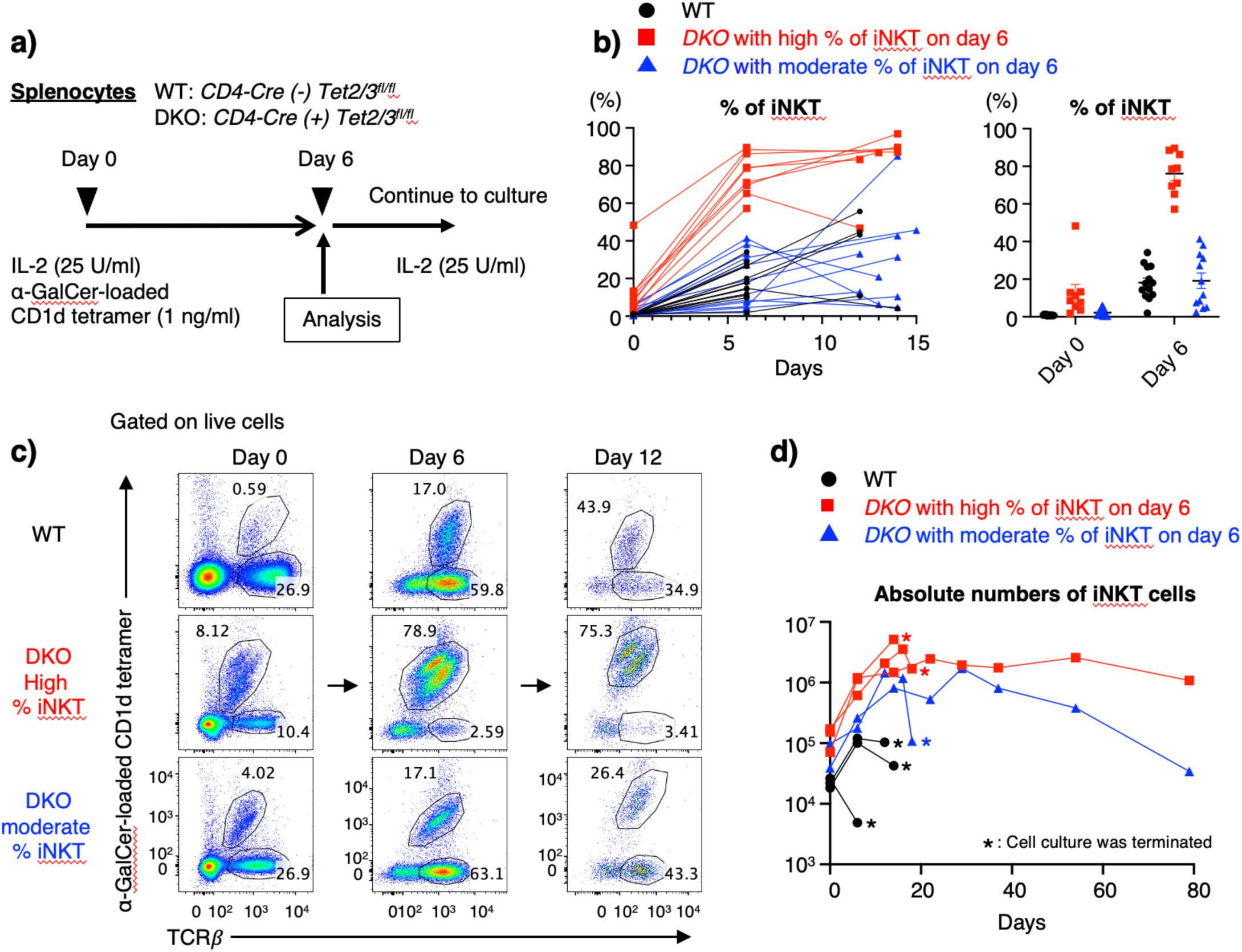
*Tet2/3*-deficient iNKT cells exhibit enhanced long-term expansion *in vitro*. (a) Experimental design. Splenocytes isolated from WT (Tet2^fl/fl^ Tet3^fl/fl^ CD4-Cre^-^) and *Tet2/3 DKO* (Tet2^fl/fl^ Tet3^fl/fl^ CD4-Cre^+^) mice were cultured in the presence of IL-2 (25 U/ml) and α-GalCer-loaded CD1d tetramer (1 ng/ml) for six days. Cells were subsequently passaged and maintained in IL-2 alone. iNKT-cell frequencies were analyzed by flow cytometry at the indicated time points. (b) (*Left*) Percentage of iNKT cells (TCRβ^+^ aGalCer-CD1d-tetramer^+^) 0-15 days after starting the culture. *Tet2/3* DKO cultures segregated into DKO-high and DKO-moderate groups based on the frequency of iNKT cells after six days of culture. (*Right*) Correlation of iNKT cells frequency between at culture initiation and frequency of iNKT cells after 6 days of cell culture. (c) Representative flow cytometry plots and quantification of iNKT-cell frequencies at day 0, day 6, and day 12. (d) Absolute numbers of WT and *Tet2/3* DKO iNKT cells during long-term culture, plotted on a log_10_ scale.

Consistent with the known responsiveness of iNKT cells to glycolipid antigen stimulation, WT iNKT cells expanded from <1% of splenocytes at the start of culture to ∼20% by day 6 (**Fig. 2b**, *black*). In contrast, *Tet2/3 DKO* cultures segregated into two distinct groups. In one group, iNKT cells expanded dramatically and exceeded 50% of viable cells by day 6 (DKO-high, *red*), whereas in the second group iNKT cells reached frequencies around 20%, comparable to those observed in WT cultures (DKO-moderate) (**Fig. 2b**, *blue*). Notably, DKO-high cultures already contained elevated frequencies of iNKT cells at the initiation of culture, averaging ∼12% of splenocytes, whereas DKO-moderate cultures contained only modestly increased frequencies (∼2%) (**Fig. 2b**, *right panel*; **Fig. 2c**). The frequencies and absolute numbers of iNKT cells at day 0 correlated strongly with their frequencies and absolute numbers after six days of culture (**Figs. 2b-d**), suggesting that although *TET* deficiency is sufficient to cause increased proliferation, some *Tet2/3*-deficient mice already harbor a subpopulation of iNKT cells with markedly enhanced proliferative potential. *Tet2/3*-deficient iNKT cells displayed clonal expansion (**Suppl. Fig. S2a**), enhanced proliferative capacity, prolonged viability, and sustained expansion for up to 79 days in culture, whereas WT cultures failed to persist beyond approximately 12-14 days (**Fig. 2d**).

To determine whether *Tet2/3 fl/fl CD4Cre* mice displayed alterations in other lymphocyte populations, we analyzed the frequencies of conventional TCRβ^+^ α-GalCer-CD1d tetramer-negative T cells within the same *Tet2/3*-deficient cultures (see **Fig. 2c**). DKO-high cultures displayed a marked reduction in tetramer-negative TCRβ^+^ cells (non-iNKT cells; **Fig. 2c** and **Suppl. Fig. S2b**). Further analysis of the TCRβ^+^ compartment revealed expansion of CD4^-^CD8^-^ double-negative T cells in *Tet2/3 DKO* cultures (**Suppl. Fig. S2c**). We presume that these cells arose from CD4^+^ cells that had lost CD4 expression due to *TET* deficiency, consistent with the requirement for continuous TET function to maintain CD4 expression in peripheral T cells through continued demethylation of certain *Cd4* enhancers (Issuree et al., 2018; Sellars et al., 2015; Suzuki et al., 2025). The CD4^-^ CD8^-^ double-negative T cells expanded from day 0 to day 6 in cultures with moderate (**Suppl. Fig. S2c,** *top panel*) as well as high (**Suppl. Fig. S2c,** *bottom panel*) frequencies of TCRβ^+^ α-GalCer-CD1d tetramer^+^ iNKT cells (**Suppl. Fig. S2c***, compare left and right panels*), but were less abundant in cultures with higher frequencies of iNKT cells at both day 0 and day 6 (**Suppl. Fig. S2c,** *top panel, compare left and right*). The expansion of CD4^-^CD8^-^ double-negative T cells may compete with (and thereby limit) iNKT-cell expansion within *Tet2/3*- deficient cultures.

Overall, the absolute number of iNKT cells recovered from both DKO-high and DKO-moderate cultures exceeded those recovered from WT cultures (**Fig. 2d**), indicating that *Tet2/3* deficiency intrinsically enhances the proliferative capacity of iNKT cells. As noted above, WT cultures could not be maintained beyond 12-14 days, whereas *TET*-deficient iNKT cultures largely remained viable and continued to proliferate for more than 2.5 months (79 days) in the presence of IL-2, even without repeated TCR stimulation with α-GalCer-loaded CD1d tetramer (**Fig. 2d**). As in many cancers that become independent of mitogenic signaling at later times (Hanahan & Weinberg, 2011), antigen stimulation was required early but the T cells could become antigen-independent— and appear to “dedifferentiate” to lose CD4—at later times. Thus, the remarkable expansion and persistence of *Tet2/3*-deficient iNKT cells observed following transplantation can be reproduced *in vitro*, establishing a tractable experimental system for investigating the mechanisms underlying *TET*-deficient iNKT cell transformation.

### Cells with chromosome 17 trisomy are selectively enriched during expansion of *Tet2/3*-deficient iNKT cells

The recurrent acquisition of chromosome 17 trisomy in *Tet2/3*-deficient iNKT-cell lymphomas raised an important question: does chromosome 17 gain arise *de novo* late during malignant progression, or does it occur as a secondary consequence of chromosome instability during early expansion of TET-deficient cells? In the latter case, we would expect to detect a pre-existing subpopulation of cells with chromosome 17 trisomy in *Tet2/3 DKO* iNKT cells from young mice prior to transfer, that would later undergo selective expansion when expanded in recipient mice. However, bulk genomic analyses of pre-transfer *Tet2/3 DKO* iNKT cells failed to detect chromosome 17 trisomy (**Fig. 1b**), suggesting that if cells with trisomies were present, they existed below the detection threshold of population-based approaches. To resolve chromosome copy-number heterogeneity, we analyzed *Tet2/3 DKO* iNKT cells before and during expansion in culture by single-cell whole-genome sequencing (scWGS) (Bakker et al., 2016).

To assess the enrichment of potentially aneuploid cells during sustained expansion, purified WT and *Tet2/3 DKO* iNKT cells were analyzed immediately after isolation (day 0) and following 6 or 13 days of culture under the expansion conditions described in **Fig. 2** (because of the poor viability of WT cultures, only *Tet2/3 DKO* cultures were analyzed at 13 days). scWGS of WT iNKT cells revealed a largely stable karyotype, with no whole-chromosome aneuploidies detected at day 0 across the two biological replicates (48 cells total; **Fig. 3a,b**). Following 6 days of culture, only two aneuploid cells were identified in WT cultures across both replicates: one cell with chromosome 12 monosomy (1/24 cells, 4.2%) and one cell with chromosome 12 trisomy and chromosome 13 monosomy (1/42 cells, 2.4%) (**Fig. 3a,b,e**).

**Figure 3.**
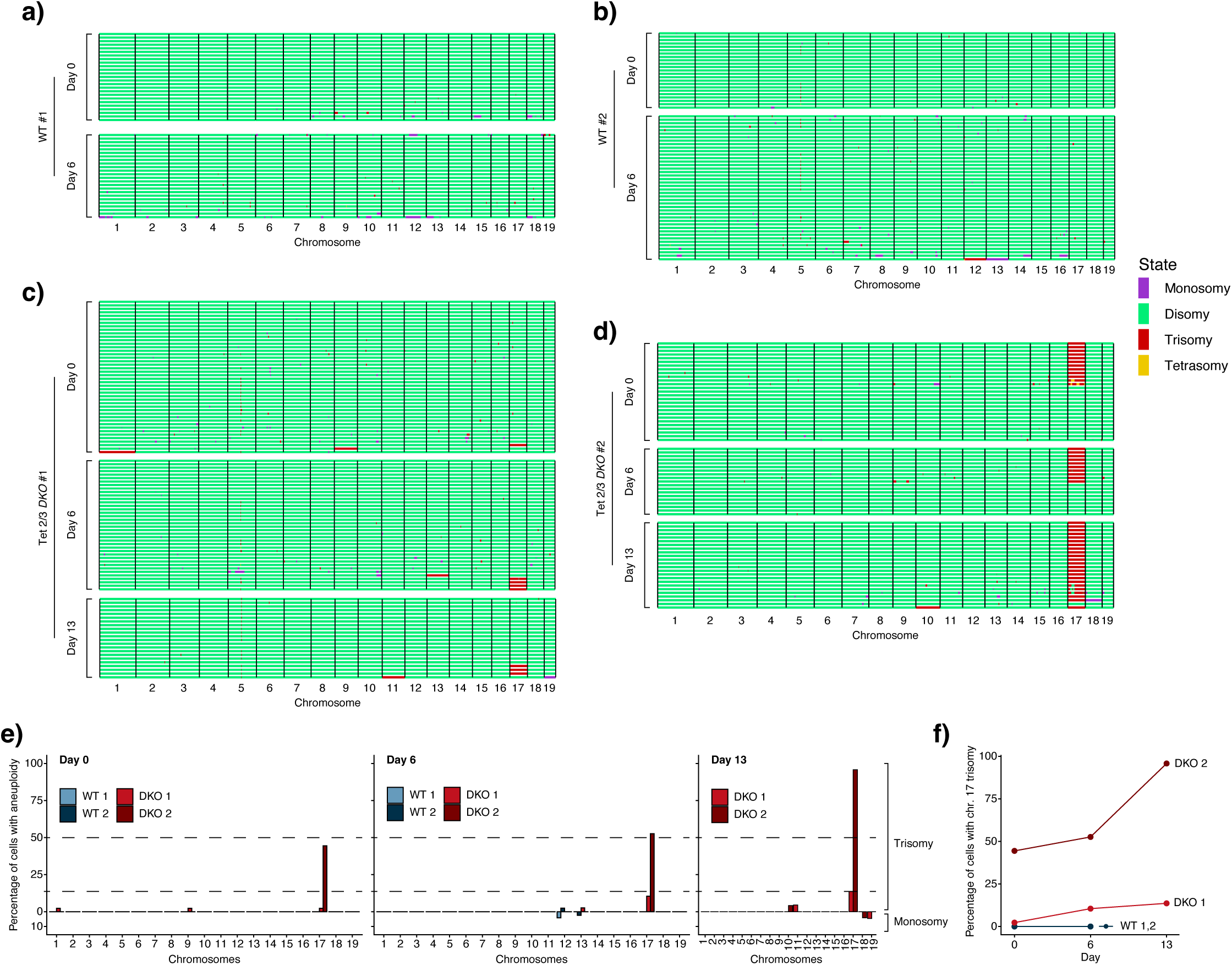
Single-cell genomic analysis reveals enrichment of chromosome 17 trisomy in *Tet2/3*-deficient iNKT cells during expansion. (**a-d**) Single-cell whole genome sequencing (scWGS) of WT (**a**, **b**) and *Tet2/3 DKO* (**c**, **d**) iNKT cells isolated from two donor mice of each genotype, analyzed at the indicated days of *in vitro* expansion. WT iNKT cells (**a**, **b**) were analyzed at days 0 and 6, whereas *Tet2/3 DKO* iNKT cells (**c**, **d**) were analyzed at days 0, 6, and 13. Each row represents an individual cell, and each column represent an individual chromosome. Colors indicate the inferred chromosome copy-number state (*see legend, right*). Cell numbers: (**a**) WT #1 day 0, n=25; WT #1 day 6, n=24; (**b**) WT #2 day 0, n=23; WT #2 day 6, n=42; (**c**) *Tet2/3 DKO* #1 day 0, n=44; *Tet2/3 DKO* #2 day 6, n=38; *Tet2/3 DKO* #2 day 13, n=22; (**d**) *Tet2/3 DKO* day 0, n=27; *Tet2/3 DKO* day 6, n=19; *Tet2/3 DKO* day 13, n=24. (e) Quantification of aneuploid cells identified by scWGS. Bars show the percentage of cells with chromosome copy-number alterations for each chromosome at the indicated days of *in vitro* expansion, displaying data from two independent mice per genotype. Bars extending above zero indicate trisomy, whereas bars extending below zero indicate monosomy. Dashed lines indicate 12.5% and 50% cell frequencies for reference. Chromosome 17 trisomy was detected at day 0 and became enriched during expansion of *Tet2/3 DKO* iNKT cells, indicating selective expansion of cells with pre-existing chromosome 17 trisomy. (f) Frequency of chromosome 17 trisomy during *in vitro* expansion of WT and *Tet2/3 DKO* iNKT cells. Each line represents an independent donor mouse (two WT and two *Tet2/3 DKO* mice), showing the percentage of cells with three copies of chromosome 17 at days 0, 6, and 13 of expansion.

In contrast, *Tet2/3 DKO* iNKT cells displayed greater karyotypic heterogeneity at day 0, and the frequency of cells with chromosome 17 trisomy increased during culture in two independent replicates (**Fig. 3c,d,f**). In one donor, chromosome 17 trisomy was detected in 1 of 44 cells at day 0 (2.3%), increased to 4 of 38 cells (10.5%) after 6 days of expansion, and to 3 of 22 cells (13.6%) by day 13 (**Fig. 3c**)—these cells expanded as a DKO- high culture. In the second donor, chromosome 17 trisomy was substantially more abundant at baseline, detected in 12 of 27 cells (44.4%) at day 0, and increased to 10 of 19 cells (52.7%) at day 6 and 23 of 24 cells (95.8%) at day 13 (**Fig. 3d**)—these cells expanded as a DKO-moderate culture. Although the abundance of the pre-existing population of cells with chromosome 17 trisomy varied markedly between animals, the direction of selection was consistent, with cells carrying chromosome 17 trisomy becoming enriched during expansion in both independent *Tet2/3 DKO* cultures (**Fig. 3c,d,f**). It was noteworthy that chromosome 17 trisomy was detected in *Tet2/3 DKO* cells at day 0 in both biological replicates by scWGS (**Fig. 3c,d**), despite being undetectable by bulk genomic analyses (**Fig. 1b**, *Young*), indicating that cells with chromosome 17 trisomy are already present as a subpopulation within *Tet2/3 DKO* mice prior to expansion.

Other whole chromosome copy-number alterations were detected sporadically in *Tet2/3 DKO* cells during culture, including trisomies involving chromosomes 10, 11, and 13, and monosomies involving chromosomes 18 and 19 (**Fig. 3c-e**). These alterations were generally detected in individual cells and did not show the recurrent enrichment observed for chromosome 17. Although copy-number alterations occurred sporadically during *in vitro* expansion of both WT and *Tet2/3 DKO* cells, the reproducible enrichment of cells with chromosome 17 trisomy across independent *Tet2/3 DKO* cultures, together with the uniform presence of chromosome 17 trisomy among expanded cells across transplantation experiments (**Fig. 1b**, *Transferred*), indicates that chromosome 17 gain is selectively favored, especially during expansion *in vivo*.

Together, these single-cell analyses demonstrate that chromosome 17 trisomy is already present in a subset of *Tet2/3*-deficient iNKT cells before expansion and becomes progressively enriched during proliferation in culture. This provides a plausible explanation for the recurrent chromosome 17 trisomy observed in independently derived transplanted iNKT-cell lymphomas (**Fig. 1a,b**) and suggests that chromosome 17 gain confers a selective fitness advantage during sustained expansion of *Tet2/3*-deficient iNKT cells. Because this enrichment can occur under defined culture conditions, these findings further suggest that the selective advantage associated with chromosome 17 gain may arise through a cell-intrinsic mechanism that does not show an absolute requirement for the *in vivo* microenvironment.

### *TET* deficiency promotes the emergence and selection of aneuploid subclones in other cell types

The enrichment of subclonal populations carrying chromosome 17 trisomy during expansion of *Tet2/3*-deficient iNKT cells raised the question of whether recurrent aneuploid selection is a specific feature of *TET*-deficient lymphoid cells or reflects a more general consequence of TET loss. To address this question, we turned to a previously established *Tet1^fl/fl^ Tet2^fl/fl^ Tet3^fl/fl^* (*Tet Tfl*) mouse embryonic stem cell (mESC) model that enables acute inducible deletion of all three *Tet* genes with a tamoxifen-inducible Cre-ERT2 transgene. In this model cell line, metaphase spreads performed 6 days after triple *Tet* gene deletion revealed substantially higher frequencies of aneuploid cells in the resulting *Tet iTKO* mESC (Georges et al., 2022). Live-cell imaging of these cultures showed increased frequencies of micronuclei, chromatin bridges, and lagging chromosomes following *Tet* deletion, indicating defects in chromosome segregation, attributed at least in part to silencing of the *Khdc3* gene (Georges et al., 2022).

*Tet Tfl* mESCs carrying the inducible *Cre-ERT2* and a *Rosa26-LSL-H2B-GFP* reporter allele were treated with 4-hydroxytamoxifen (4-OHT) to induce simultaneous deletion of all three *Tet* genes and activation of the H2B- GFP reporter (**Fig. 4a**). *Tet* gene deletion in these cells is highly efficient as previously reported (Georges et al., 2022). Following three days of 4-OHT treatment and three additional days of culture after drug withdrawal, *Tet*- inducible triple-knockout (*Tet iTKO*) cells and parallel untreated *Tet* triple-floxed (*Tet Tfl*) control cells were analyzed by scWGS (**Fig. 4a**). Two independently derived mESC clones were analyzed in parallel, with each clone generating matched *Tet Tfl* and *Tet iTKO* populations.

**Figure 4.**
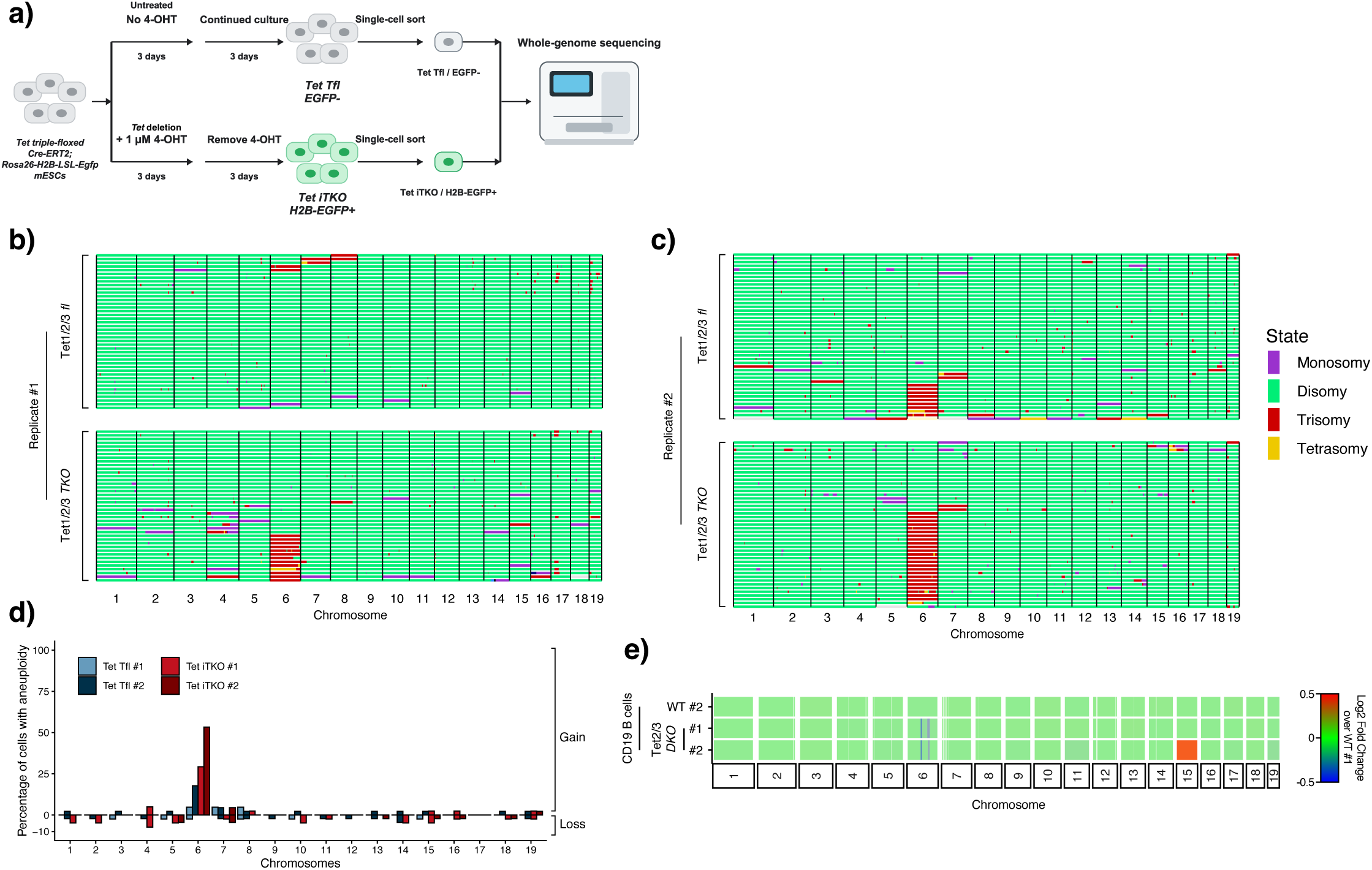
*TET* deficiency promotes recurrent chromosome gains across cell types. (**a**) Experimental design. Tet1^fl/fl^ Tet2^fl/fl^ Tet3^fl/fl^ mESCs carrying Cre-ERT2 and Rosa26-LSL-H2B-GFP alleles were treated with 1 μM 4-hydroxytamoxifen (4-OHT) for three days to induce deletion of all three *Tet* genes and activation of H2B-GFP expression (*Tet iTKO* cells). Control cells were not treated with 4-OHT (*Tet Tfl* cells). Following 4-OHT withdrawal, cells were cultured for an additional three days before analysis. Two independently derived ESC clones were analyzed in parallel, yielding paired *Tet Tfl* and *Tet iTKO* samples (Tet Tfl #1, Tet iTKO #1, and Tet Tfl #2, Tet iTKO #2). (**b, c**) Single-cell whole genome sequencing (scWGS) of two independently derived *Tet Tfl* and *Tet iTKO* mESC clone pairs (**b**, clone #1; **c**, clone #2). Each row represents an individual cell, and each column represents an individual chromosome. Colors indicate the inferred chromosome copy-number state (*see legend, right*). *Tet*- deficient cells display increased karyotypic heterogeneity and recurrent chromosome 6 trisomy relative to control cells. Cell numbers: (**b**) *Tet Tfl* #1, n=42; *Tet iTKO* #1, n=41; (**c**) *Tet Tfl* #2, n=45; *Tet iTKO* #2, n=45. (d) Quantification of aneuploid cells identified by scWGS. Bars show the percentage of cells with chromosome copy-number alterations for each chromosome in the *Tet Tfl* and *Tet iTKO* mESC samples. Bars extending above zero indicate chromosome gains, whereas bars extending below zero indicate chromosome losses. *Tet-*deficient mESC clones showed increased frequencies of aneuploid cells, with chromosome 6 gain representing the most prominent recurrent copy-number alteration. (e) Genome-wide copy-number profiles of WT and *Tet2/3 DKO* CD19^+^ B cells, inferred from whole-genome bisulfite sequencing (WGBS) datasets previously published in (Shukla et al., 2022). Each point represents a 100- kb genomic window. Red indicates copy-number gain. Chromosome 15 gain was observed in one of two independent *Tet2/3 DKO* samples.

scWGS revealed increased karyotypic heterogeneity in *Tet iTKO* mESCs compared to control *Tet Tfl* cells, with individual cells harboring diverse whole-chromosome gains and losses (**Fig. 4b-d**). Chromosome 6 trisomy was detected at baseline in control cultures and increased following *Tet* deletion in both independently derived clones, although its abundance and degree of enrichment varied between clones (**Fig. 4b,c**). Chromosome 6 trisomy is a recurrent chromosomal abnormality observed in mESC during culture, and mESCs carrying trisomy 6 exhibit increased proliferation compared with euploid mESCs (Montilla-Rojo et al., 2025; Zhang et al., 2016). Consistent with these observations, chromosome 6 trisomy represented the most prominent recurrent chromosome alteration across the two independently derived *Tet iTKO* mESC populations, whereas additional chromosome gains and losses occurred sporadically in individual cells (**Fig. 4d**). Together, these findings indicate that *TET* deficiency promotes the emergence of diverse aneuploid states, with chromosome 6 gain preferentially enriched during expansion of *TET*-deficient mESCs.

The enrichment of cells with chromosome 6 trisomy in *Tet iTKO* mESC cultures parallels the selective enrichment of cells with chromosome 17 trisomy during *Tet2/3*-deficient iNKT cell expansion (**Fig. 1b**; **Fig. 3**). Notably, however, the chromosome preferentially enriched is different between the mESC and T cells. To determine whether aneuploid populations also emerge in other primary hematopoietic cells following *TET* deficiency, we inferred chromosome copy-number alterations in B cells by analyzing previously generated WGBS data from WT and *Tet2/3 DKO* CD19^+^ B cells isolated from a mouse model of *Tet2/3*-deficient B-cell lymphoma (Shukla et al., 2022). One of two independent *Tet2/3 DKO* B-cell lymphoma samples displayed chromosome 15 trisomy (**Fig. 4e**), providing additional evidence that chromosome gains can emerge in distinct cellular contexts following *TET* deficiency. Notably, these B cell populations were not subjected to the transplantation and expansion scheme used for the *Tet2/3*-deficient iNKT cell experiments.

Across these distinct cellular contexts, different chromosomes were preferentially enriched: chromosome 17 in iNKT cells, chromosome 15 in B cells, and chromosome 6 in mESC. Thus, *TET* deficiency promotes the emergence of diverse aneuploid states, with different chromosomes preferentially gained in different cell types. This observation argues against a chromosome-specific effect of *TET* loss and instead suggests that *TET* deficiency creates a permissive environment for the emergence of aneuploid subclones, while the particular chromosome that is amplified is determined by cell-type-specific selective pressures. Our findings extend the observation of recurrent aneuploid selection beyond the lymphoid lineage, and support a model in which disruption of TET-mediated epigenetic regulation promotes the generation and expansion of cryptic aneuploid subpopulations across diverse cellular contexts.

### Loss of *Dnmt3a* is associated with recurrent chromosome gains in lymphoid malignancies

The recurrent selection of chromosome 17 trisomy in *Tet2/3*-deficient iNKT cells, together with the appearance of aneuploid populations in other *TET*-deficient models, raised the question of whether recurrent chromosome gains represent a broader consequence of perturbing DNA methylation and demethylation pathways. To address this question, we analyzed publicly available WGBS datasets from mouse models of *Dnmt3a*- and *Dnmt3b*- deficient lymphoid malignancies, including chronic lymphocytic leukemia (CLL), peripheral T-cell lymphoma (PTCL), and MYC-induced T-cell lymphoma (Haney et al., 2016; Lopusna, Nowialis, Opavska, Abraham, Riva, Haney, et al., 2021; Lopusna, Nowialis, Opavska, Abraham, Riva, & Opavsky, 2021). *Dnmt3a* and *Dnmt3b* encode the two *de novo* DNA methyltransferases, with important roles in hematopoietic development and lymphoid differentiation (Challen et al., 2011; Challen et al., 2014).

Copy-number profiles inferred from these datasets revealed recurrent chromosome gains in *Dnmt3a*-deficient lymphoid malignancies (**Fig. 5**). Chromosome 17 trisomy was detected in two of four *Dnmt3a*-deficient samples, including one of two CLL samples and one of two PTCL samples. Chromosome 15 gain was more frequent, occurring in three of four *Dnmt3a*-deficient samples, including both CLL samples and one of two PTCL samples. Chromosome 15 gain was also detected in one of four *Dnmt3b*-deficient samples, specifically in one of two MYC-induced T-cell lymphoma samples (**Fig. 5**). Thus, disruption of *de novo* DNA methyltransferase function is associated with recurrent chromosome gains in lymphoid malignancies, although the frequency and identity of these alterations vary between models.

**Figure 5.**
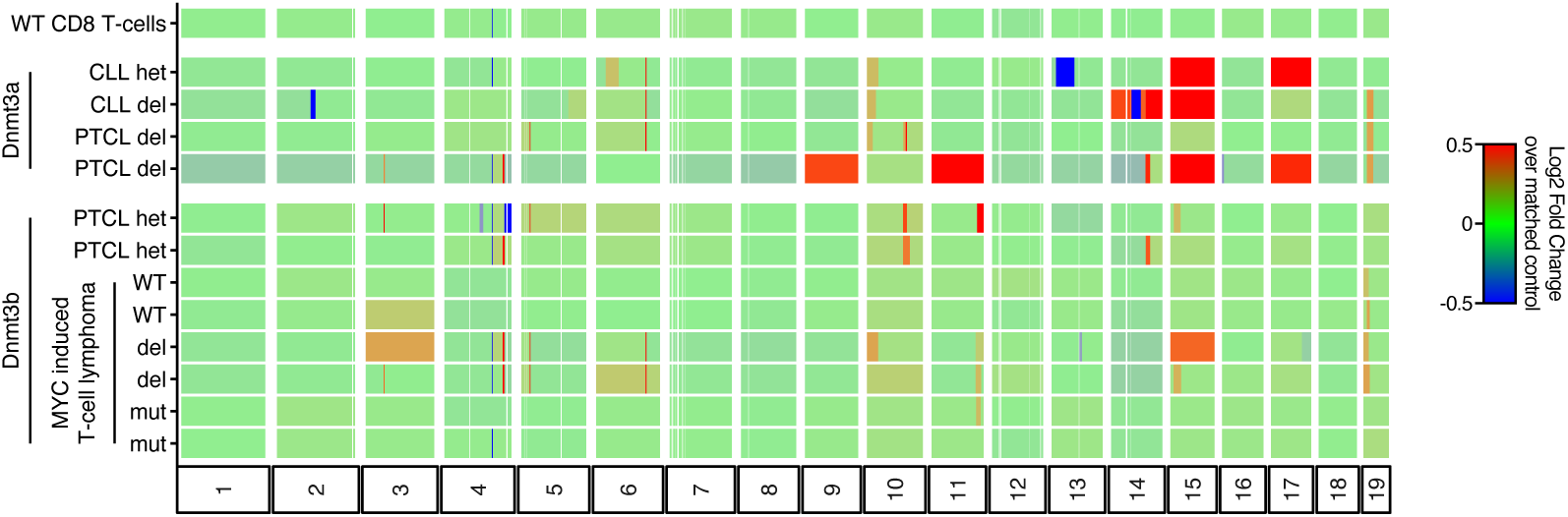
Recurrent chromosome gains associated with *Dnmt3a* and *Dnmt3b* deficiency in mouse models of lymphoid malignancies. Genome-wide copy-number profiles of *Dnmt3a*-deficient (*het*: heterozygous knockout; *del*: homozygous knockout) and *Dnmt3b*-deficient or catalytically inactive mutant (*mut*) lymphoid malignancies, including chronic lymphocytic leukemia (CLL), peripheral T-cell lymphoma (PTCL), and MYC-induced T cell lymphoma, inferred from whole-genome bisulfite sequencing (WGBS) datasets previously published in (Haney et al., 2016; Lopusna, Nowialis, Opavska, Abraham, Riva, Haney, et al., 2021; Lopusna, Nowialis, Opavska, Abraham, Riva, & Opavsky, 2021)). Each point represents a 100-kb genomic window. Red indicates copy-number gain and blue indicates copy-number loss. Chromosome 17 trisomy was observed in two of four *Dnmt3a*-deficient samples, including one of two CLL samples and one of two PTCL samples. Recurrent chromosome 15 gain was observed across independently derived *Dnmt3a*- and *Dnmt3b*-deficient models, occurring in three of four *Dnmt3a*-deficient samples (two of two CLL and one of two PTCL) and one of four *Dnmt3b*-deficient samples (in one of two MYC- induced T cell lymphoma samples).

The recurrence of chromosome 17 trisomy in two of four *Dnmt3a*-deficient lymphoid malignancies was particularly notable given its recurrent enrichment in *Tet2/3*-deficient iNKT-cell lymphoma. Chromosome 15 gain was also observed across *Dnmt3a*- and *Dnmt3b*-deficient models and has been observed in other *TET*-deficient lymphoid populations, including *Tet2/3*-deficient B-cell lymphomas (this study) and expanded *Tet2/3*-deficient iNKT cells following repeated transfer (Gioulbasani et al., 2024). Together, these observations suggest that opposing perturbations of DNA methylation and demethylation pathways can converge on similar patterns of chromosome gain.

Our observations extend the association between disruption of DNA methylation homeostasis and evolution of recurrent aneuploidies beyond *TET* deficiency, and suggest that dysregulation of DNA methylation and demethylation pathways creates conditions permissive for the emergence and selection of advantageous aneuploid subclones during lymphoid expansion. Our data support a model in which the specific chromosomal alterations that become enriched are shaped by the underlying genetic perturbation, cellular context, lineage identify, and selective pressures from cytokines or surrounding cells.

## Discussion

In this study, we establish that recurrent aneuploidies are a prominent feature of cellular expansion following *TET* deficiency. Across complementary *in vivo* mouse models and in cultured cells, our data show that loss of TET function generates a reservoir of karyotypic diversity from which advantageous chromosomal configurations can emerge during extended proliferation. The clearest example was chromosome 17 trisomy in *Tet2/3*-deficient mouse iNKT-cell lymphomas, which remarkably was detected in ten of ten independently derived tumors following transplantation, with other chromosome gains and losses varying between tumors (**Fig. 1b**). Consistent with this finding, expanded non-iNKT, non-Treg T cells from the same *Tet2/3 fl (CD4Cre*) strain of mice also showed recurrent enrichment of chromosome 17 following serial transplantation, together with variable aneuploidies involving other chromosomes (Gioulbasani et al., 2024).

Single-cell analysis of *Tet2/*3 *DKO* mouse iNKT cells further revealed that cells carrying chromosome 17 trisomy were already present as a rare subpopulation before expansion (**Fig. 3c-d**, day 0). Other single cells carried different aneuploidies that varied between cells, but the subpopulation with chromosome 17 trisomy became progressively enriched during cellular expansion following transplantation (**Fig. 1b**) or subsequent culture (**Fig. 3c-f**). Our findings distinguish two related but mechanistically distinct processes: *TET* deficiency quickly increases karyotypic diversity, generating a reservoir of aneuploid subpopulations, whereas extended cellular proliferation provides the opportunity for specific aneuploid subclones to undergo positive selection.

The rapid detection of increased aneuploidy and copy-number alterations in *TET*-deficient cells—pre-transfer iNKT cells from young *Tet2/3*-deficient mice (**Fig. 3c,d**) or mESCs six days after acute *Tet* deletion (**Fig. 4b,c**)— suggests that *TET* deficiency may directly or indirectly promote aneuploidy through defects in chromosome segregation arising from altered epigenetic regulation. Indeed, recent studies have linked defects in DNA methylation to centromeric and pericentromeric dysfunction: briefly, targeting a fusion protein of the TET1 catalytic domain with the CENP-B DNA binding domain to promote DNA demethylation at α-satellite regions compromised centromere integrity and resulted in chromosomal instability (Salinas-Luypaert et al., 2025). In another study, hypermethylation of the *Khdc3* promoter and its associated transcriptional downregulation, which occurs in mESC with triple *Tet* deficiency, was invoked to explain the increased frequencies of lagging chromosomes, chromatin bridges, micronuclei, and aneuploidy observed within 6 days of deletion of all three *Tet* genes in mESCs (Georges et al., 2022). An alternative hypothesis, not mutually exclusive with promoter hypermethylation, involves the hypomethylation within heterochromatic regions that we previously described in diverse proliferating *TET*-deficient cells (López-Moyado et al., 2026; López-Moyado et al., 2019). If TET-dependent heterochromatin hypomethylation extended to centromeric or pericentromeric regions, it might contribute to chromosome segregation defects and aneuploidy. Regardless of which mechanism predominates, our findings support a model in which loss of *TET* activity promotes chromosome segregation errors, thereby increasing the generation of aneuploid cells upon which selection can subsequently act. Such a mechanism could help explain the genomic instability, DNA damage, and oncogenic transformation observed in multiple cancers, not only blood cancers with *TET2* deficiency but also a variety of solid cancers with mutations or silencing of *TET1, TET2* or *TET3* (Bray et al., 2021).

The second process, the selection of specific aneuploid states required additional proliferation. Specifically, chromosome 17 trisomy became prominent only after *Tet2/3*-deficient iNKT cells, which had already expanded in young *Tet2/3 DKO (CD4Cre)* mice, were transferred into immunocompetent recipient mice where they underwent further expansion (López-Moyado et al., 2019; Tsagaratou et al., 2017). Supporting this conclusion, *TET*-deficient mESC displayed greater karyotypic diversity than control *Tet Tfl* cells cultured for the same number of days, yet we observed that only chromosome 6 trisomy was recurrently enriched in *Tet iTKO* cells (**Fig. 4b-d**). Thus, the recurrent chromosome gains observed in *TET*-deficient cells are unlikely to reflect solely the stochastic accumulation of chromosome segregation errors; rather, continued proliferation permits the selection of particular aneuploidies that confer a fitness advantage in a given cellular context. Similar conclusions have been drawn from more elaborate cancer models, where an early predisposition to aneuploidy is subject to *in vivo* selection and gives rise to specific aneuploid states (Foijer et al., 2017; Foijer et al., 2014; Shoshani et al., 2021; Trakala et al., 2021).

Collectively, these findings suggest that *TET* deficiency increases aneuploidy and karyotypic diversity, whereas continued proliferation is required to determine which aneuploid states become enriched through selection. The reproducible enrichment of chromosome 17 trisomy in *Tet2/3 DKO TET*-deficient iNKT cells indicates that increased chromosome 17 dosage provides a strong proliferative advantage during expansion of these cells. Genes located on chromosome 17 exhibited increased expression proportional to chromosome dosage and were enriched for functions related to RNA processing, ribosome biogenesis, chromosome segregation, and mitochondrial activity, processes that may collectively enhance proliferative fitness. Our data from cells cultured under minimal defined stimulation suggests that selection for chromosome 17 trisomy is at least partly cell-intrinsic, and that it requires both *TET* deficiency and TCR/IL-2 stimulation. Further studies will be necessary to identify the specific genes on chromosome 17 that, either individually or together, confer this strong proliferative advantage.

Similarly, the preferential enrichment of chromosome 6 trisomy in *TET*-deficient mESCs is consistent with a growth advantage of this karyotype in mESCs, as chromosome 6 gain is a well-documented recurrent feature of mESCs during prolonged culture, and mESCs carrying chromosome 6 trisomy exhibit enhanced proliferation and reduced dependence on exogenous growth factors for self-renewal (Montilla-Rojo et al., 2025; Zhang et al., 2016). Thus, selection for different chromosomes in *TET*-deficient iNKT cells and mESCs further suggests that the fitness advantage conferred by aneuploidy is shaped by cellular context rather than by an intrinsic property of a single chromosome.

Our conclusions extend to other regulators of DNA cytosine methylation. Mouse models of *Dnmt3a*-deficient lymphoid malignancies displayed recurrent chromosome 17 trisomy and, even more frequently, chromosome 15 trisomy (**Fig. 5**). Chromosome 15 trisomy was also observed in *Dnmt3b*-deficient T-cell lymphoma (**Fig. 5**) and *Tet2/3*-deficient B-cell lymphoma (**Fig. 4e**). Mice carrying a hypomorphic *Dnmt1* allele that causes widespread DNA hypomethylation also developed aggressive T-cell lymphomas with a high frequency of chromosome 15 trisomy (Gaudet et al., 2003). Thus, recurrent selection for specific aneuploidies can arise following disruption of distinct components of the DNA methylation machinery. The convergence of *TET* and *DNMT* deficiency in selecting for recurrent gains of overlapping chromosomes is particularly notable because these enzymes have opposing enzymatic activities in the regulation of DNA methylation. TET proteins promote DNA demethylation whereas DNMTs catalyze DNA methylation, yet disruption of either pathway appears capable of generating epigenetic changes that are permissive for the selection of specific chromosomal gains.

The recurrence of particular chromosome gains across independent experimental systems suggests that some aneuploid states may provide reproducible fitness advantages during lymphoid expansion or tumor evolution. Chromosome 17 trisomy and chromosome 15 trisomy have been reported in the IL-15 transgenic mouse model of aggressive NK/T-cell large granular lymphocyte leukemia (Yokohama et al., 2010), a disease that shares several biological features with the proliferative lymphoid disorders studied here. Similarly, karyotyping of iNKT-cell lymphomas from *Id2/Id3*-deficient mice revealed chromosome 17 trisomy in one of two tumors analyzed (Li et al., 2017). The recurrent trisomies of chromosomes 15 and 17 across genetically and experimentally distinct models of lymphoid neoplasia emphasizes that specific chromosome gains are repeatedly favored during lymphoid tumorigenesis, rather than arising merely as stochastic byproducts of genome stability. Noteworthy, the *Myc* gene is located on mouse chromosome 15, *MYC* amplification is well established in human cancers (Dhanasekaran et al., 2022), and previous studies have implicated MYC in the recurrent selection of chromosome 15 trisomy in p53-deficient murine T-cell malignancies (Shoshani et al., 2021; Trakala et al., 2021). However, other genes located on chromosomes 15 and 17 may also be involved.

In conclusion, aneuploidy may represent an early step in the evolution of malignancies driven by *TET2* or *DNMT3A* loss, with potential implications for lymphoid malignancies associated with altered DNA methylation regulation, and dysregulation of DNA methylation may influence not only the generation of aneuploidy, but also which chromosomal alterations are ultimately favored during tumor development. For instance, among blood cancers, both DLBCL and PTCL show a greater propensity to aneuploidies than T-ALL, B-ALL, AML, MDS, CMML (**Suppl Fig. 3**). Although *TET2* mutations have been reported in only a subset of PTCL (Suehara et al., 2025) and DLBCL (Schmitz et al., 2018), their presence defines biologically distinct disease subsets in which the relationship between DNA methylation dysregulation, aneuploidy, and disease progression merit further investigation. Similar considerations apply to *DNMT3A* mutations and alterations in other components of the DNA methylation-demethylation machinery, including secondary regulators (such as kinases, phosphatases or other enzymes that deposit or remove post-translational modifications) whose alteration could indirectly perturb TET or DNMT activity without mutations in the enzymes themselves.

## Methods

### Mice

*Tet2^fl/fl^Tet3^fl/fl^ CD4-Cre* mice were generated in our laboratory using B6/Taconic background “artemis” ESCs. All mice were on the B6 background and maintained in a specific pathogen-free animal facility in the La Jolla Institute for Immunology. 4-6 weeks old mice were used for each experiment since *Tet2^fl/fl^Tet3^fl/fl^ CD4-Cre* mice usually die within 8 weeks (Tsagaratou et al., 2017). All animal procedures were reviewed and approved by the Institutional Animal Care and Use Committee of the La Jolla Institute for Immunology and were conducted in accordance with institutional guidelines. Euthanasia methods complied with the AVMA Guidelines for the Euthanasia of Animals.

### Cell preparation, flow cytometry and cell sorting

Single-cell suspensions were prepared from spleen or cultured cells for staining or cell sorting. Single-cell suspensions were stained with anti-mouse antibodies or CD1d tetramer against the following (Clone name, conjugated fluorescence, dilution, manufacturer and catalog number shown in brackets): TCRβ (H57-597, BV421, 1:200, Biolegend, #109230), CD4 (RM4-5, BV605, 1:200, Biolegend, #100548), CD8 (53-6.7, PE-Cy7, 1:200, Biolegend,#100722), PBS-57 (Structural analogs of α- GarCer)-loaded CD1d tetramer (PE, 1:200, NIH Tetramer Core Facility (NIH Contract 75N93020D00005 and RRID:SCR_026557)). Flow cytometry analysis was performed using Celesta, LSR-II and Fortessa (BD Biosciences). Cell sorting was performed by the LJI flow cytometry core, using FACS ARIA-II (BD Biosciences) flow cytometers.

### *In vitro* culture for iNKT cells

Splenocytes isolated from WT (*Tet2^fl/fl^ Tet3^fl/fl^ CD4-Cre^-^*) or *Tet2/3 DKO* (*Tet2^fl/fl^ Tet3^fl/fl^ CD4-Cre^+^*) mice were cultured in the presence of IL-2 (25 U/ml) and PBS-57 (Structural analogs of α- GarCer)-loaded CD1d tetramer (1 ng/ml) for six days. Cells were subsequently passaged and maintained in IL- 2 (25 U/ml) alone.

### TCR repertoire analysis of cultured iNKT cells

For TCR repertoire analysis, cells originating from three *Tet2/3* wild-type (Tet2^fl/fl^ Tet3^fl/fl^ CD4-Cre^-^) and four *Tet2/3 DKO* (Tet2^fl/fl^ Tet3^fl/fl^ CD4-Cre^+^) mice were analyzed. Live cells from splenocytes were collected without prior cell sorting at days 0, 6, and 12 of culture under iNKT cell expansion conditions. Total RNA was extracted from the collected cells, and cDNA was synthesized. Subsequently, TCRβ-chain amplicons were generated by PCR using a panel of Vβ-specific forward primers— MuBV8.1N (5’-GGC TGA TCC ATT ACT CAT ATG TC-3’), MuBV8.2N (5’-TCA TAT GGT GCT GGC AGC ACT G-3’), MuBV7 (5’-TAC AGG GTC TCA CGG AAG AAG C-3’), MuBV2 (5’-TCA CTG ATA CGG AGC TGA GGC-3’), MuBV8.3 (5’-TGC TGG CAA CCT TCG AAT AGG A-3’), MuBV9 (5’-TCT CTC TAC ATT GGC TCT GCA GGC-3’)—and the TCRβ constant-region reverse primer MuTCB3C (5’-AAG CAC ACG AGG GTA GCC T-3’). The resulting amplicons were prepared for sequencing using the NEBNext Ultra II DNA Library Prep Kit (NEB) and sequenced on an Illumina Miseq platform.

Sequence reads were analyzed using MiXCR (Bolotin et al., 2015) to identify TCRβ clonotypes and extract CDR3 sequences. TCR repertoire abundance was quantified based on the frequency of individual clonotypes among the sequencing reads.

### Mouse embryonic stem cell culture

Previously described *Tet* triple-floxed mouse embryonic stem cells (*Tet* Tfl mESCs), carrying floxed *Tet1*, *Tet2* and *Tet3* alleles, Cre-ERT2 and a Rosa26-LSL-H2B-EGFP reporter, were used in this study (Georges et al., 2022). Cells were maintained on mitomycin C-treated mouse embryonic fibroblast (MEF) feeders in KnockOut DMEM (Thermo Fisher Scientific, 10829018) supplemented with 15% KnockOut Serum Replacement (KOSR; Thermo Fisher Scientific, 10828028), leukemia inhibitory factor (LIF), 2 mM L-glutamine, 1× MEM nonessential amino acids and 50 μM β-mercaptoethanol.

### Inducible deletion of *Tet* genes in *Tet Tfl* mESCs

As previously described (Georges et al., 2022), to induce simultaneous deletion of *Tet1*, *Tet2* and *Tet3*, *Tet* Tfl mESCs were treated with 1 μM 4-hydroxytamoxifen (4-OHT; Tocris, 3412) for 3 days, followed by culture in 4-OHT-free medium for an additional 3 days before analysis. Cre-mediated recombination generated inducible *Tet* triple-knockout (*Tet* iTKO) mESCs and concomitantly activated H2B-EGFP expression. For each of two independently derived *Tet* Tfl clones, cells cultured without 4-OHT served as matched controls, yielding four experimental groups: *Tet* Tfl #1, *Tet* iTKO #1, *Tet* Tfl #2 and *Tet* iTKO #2.

### Copy number alteration analysis of bulk samples

A subset of 10 million reads was generated for each sample. For paired-end sequencing data, only one read file was used to generate the subset. Reads were trimmed using Trim Galore (v0.6.10) and aligned to the mm10 reference genome using either Bowtie (v2.1.0) for whole-genome sequencing (WGS) and Hi-C datasets, or Bismark (v0.25.1) for whole-genome bisulfite sequencing (WGBS) datasets, with default parameters.

The resulting BAM files where subsequently analyzed in R (v4.4.1), using the Bioconductor package QDNAseq (v1.40.0) to estimate copy number alterations. A bin size of 100 kb was used. For the *applyFilters* function, the *residual*, *blacklist*, *mappability*, and *bases* parameters were set to *TRUE*. For each batch, a WT sample was used as a reference to normalize the remaining samples using the *compareToReference* function. Log_2_- transformed ratio values for the segments generated using the *segmentBins* function were plotted, using parameters *transformFun=“sqrt”* and *smoothBy = 5*.

### RNA-seq analysis

RNA-seq reads were trimmed using Trim Galore (v0.6.10) with default parameters. Trimmed reads were aligned to the mouse mm10 reference genome using STAR (v2.7.11a). Read counts were subsequently assigned to genes using featureCounts (v1.6.5) based on the NCBI mm10 RefSeq gene annotation. Downstream analyses were performed in R (v4.4.1) using Bioconductor packages. Differential gene expression analysis was performed using DESeq2 (v1.44.0), and Gene ontology (GO) enrichment analysis was performed using clusterProfiler (v4.12.6).

### Single-cell whole-genome sequencing to detect aneuploidies

Single-cell whole-genome sequencing (scWGS) was performed at iPsomics (the Netherlands) using in-house developed protocols and standard operating procedures (SOPs). Briefly, single nuclei were isolated by fluorescence-activated cell sorting (FACS) into multiwell plates. Individual nuclei were processed for whole-genome library preparation, including cell lysis, DNA fragmentation, adapter ligation with unique dual-index barcodes, and library amplification. Libraries were purified, quality controlled, and sequenced on an Illumina sequencing platform. Sequencing data were demultiplexed based on sample-specific barcodes and converted to FASTQ files. Reads were aligned to the reference genome to generate BAM files, which were subsequently analyzed for copy number alterations and aneuploidy using the AneuFinder pipeline (Bakker et al., 2016).

## Data availability

Previously published sequencing datasets used in this study are available under Gene Expression Omnibus (GEO) accession numbers GSE66834, GSE134396 and PRJNA555063 (*Tet2/3*-deficient and control iNKT cell datasets); GSE161463 (*Tet2/3*-deficient and control B cell datasets); and GSE78146, GSE154270 and GSE154451 (*Dnmt3a*-deficient and *Dnmt3b*-deficient or mutant murine lymphoid malignancy datasets). Single-cell whole-genome sequencing (scWGS) data produced as part of this study are in the process of being submitted to the European Nucleotide Archive (ENA) database and will be made publicly available prior to publication.

## Acknowledgements

We thank the NIH Tetramer Core Facility (NIH Contract 75N93020D00005 and RRID:SCR_026557) for providing mouse PBS-57-loaded CD1d tetramer PE labeled and PBS-57-loaded CD1d tetramer unlabeled. We also thank the Department of Laboratory Animal Care and the animal facility at La Jolla Institute for Immunology (LJI) for excellent support; and C. Kim, S. Alarcon, and colleagues of the La Jolla Institute Flow Cytometry and Next-Generation Sequencing (RRID:SCR_023107) Core Facilities for help with cell sorting and next-generation sequencing, respectively. The FACSAria II Cell Sorter (S10RR027366), the NovaSeq 6000 (S10OD025052) were acquired through the Shared Instrumentation Grant Program. We also thank Drs. Hyungseok Seo and Atsushi Onodera for their contributions to establishing the *in vitro* iNKT cell culture methods and optimizing culture conditions during their time in the Rao laboratory at LJI. Work in the Rao lab was supported by NIH grants R01 AI128589 and R35 CA210043 to A.R. Work in the Foijer lab is supported by a Dutch Science Council (ZonMw/NWO) Vici grant (09150182210049) to F.F. iPsomics is supported by a Dutch Science Council (ZonMw/NWO) PharmaNL grant to F.F.

## Author contributions

I.F.L.-M., K.S., and A.R. designed research and analyzed data; K.S. performed all experiments involving iNKT cells; X.L. performed all experiments involving mESCs; I.F.L.-M., and L.H.-E. performed bioinformatic analyses; D.K. and R.W. performed the scWGS data acquisition and analysis under the supervision of F.F.; F.F. and A.R. obtained funding; and I.F.L.-M. and A.R. wrote the manuscript with input from all the authors.

## Supplementary materials

Supplementary Figures S1-S3

Supplementary Tables S1-S3

**Supplementary Table S1.** Differentially expressed chromosome 17 genes in *Tet2/3 DKO* iNKT cells recovered after transplantation relative to WT iNKT cells.

**Supplementary Table S2.** Gene ontology (GO) categories and associated genes enriched among upregulated chromosome 17 genes in transplanted *Tet2/3 DKO* iNKT cells.

**Supplementary Table S3.** Expression of major histocompatibility complex (MHC) genes located on chromosome 17 in *Tet2/3 DKO* iNKT cells relative to WT iNKT cells.

**Supplementary Figure S1.**
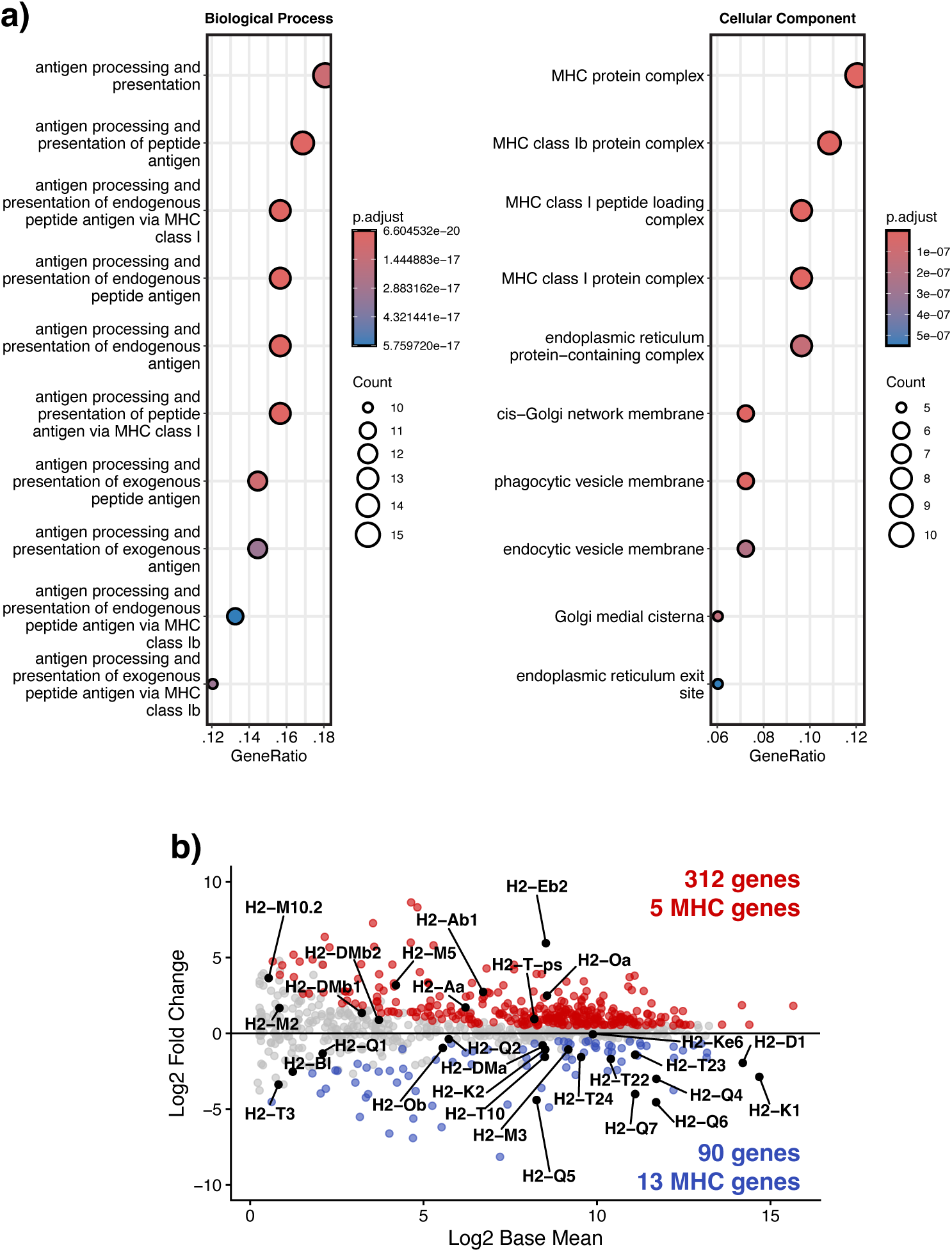
Downregulation of major histocompatibility complex genes located on mouse chromosome 17 in *Tet2/3 DKO* iNKT cells. (a) Gene Ontology enrichment analysis of chromosome 17 genes downregulated (log_2_FC<0 and *P*-adjusted value <0.05 using the Benjamini–Hochberg test) in transplanted *Tet2/3 DKO* iNKT cells relative to WT iNKT cells isolated from the spleen. Enriched categories include antigen processing and presentation. (b) MA plot of RNA-seq data showing differential gene expression of chromosome 17 genes in *Tet2/3 DKO* iNKT cells recovered after transplantation relative to WT iNKT cells. Genes meeting the criteria for differential expression (|log_2_FC|>0.5 and *P*-adjusted value <0.05 using the Benjamini–Hochberg test) are indicated in red (upregulated) or blue (downregulated). Major histocompatibility complex (MHC) genes located on chromosome 17 are highlighted by name.

**Supplementary Figure S2.**
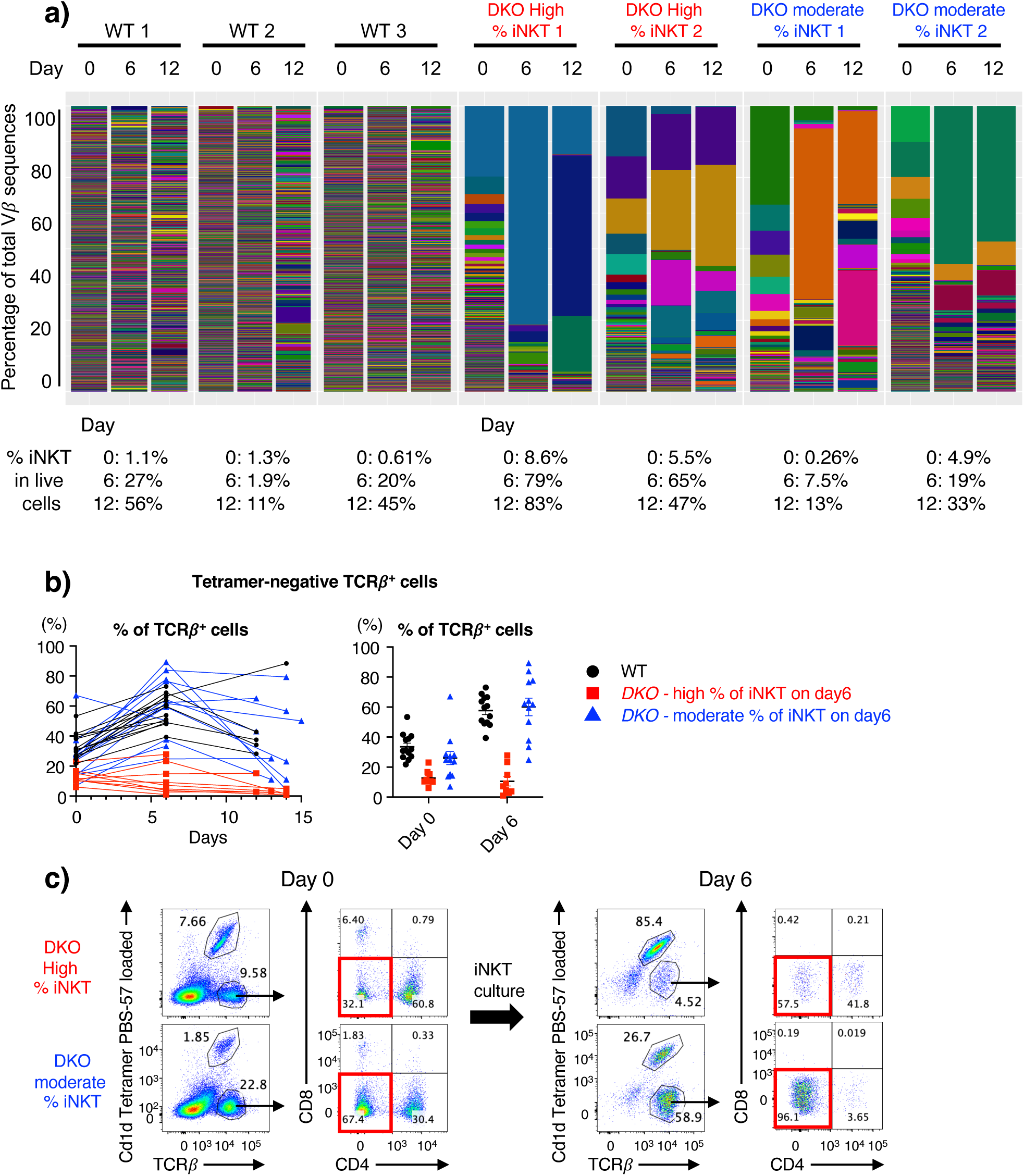
TCRβ clonality and characterization of non-iNKT populations in *Tet2/3*- deficient cell cultures. (a) TCRβ-chain clonality analysis of *Tet2/3 DKO* splenocytes in culture by amplicon sequencing. Live, unsorted splenocytes from three WT (Tet2^fl/fl^ Tet3^fl/fl^ CD4-Cre^-^) and four *Tet2/3 DKO* (Tet2^fl/fl^ Tet3^fl/fl^ CD4-Cre^+^) mice were collected at days 0, 6, and 12 of culture under iNKT cell expansion conditions. TCRβ-chain sequences were amplified from cDNA and analyzed by amplicon sequencing to assess clonality changes during culture. Each color represents one particular TCRβ sequence. (b) Quantification of TCRβ^+^ aGalCer-CD1d-tetramer^-^ conventional T cells in WT, DKO-moderate, and DKO-high cultures. (c) Frequency of double-negative CD4^-^CD8^-^ cells (ex-CD4 T cells) in TCRβ^+^ aGalCer-CD1d-tetramer^-^ T cells at day 0 and day 6 of culture.

**Supplementary Figure S3.**
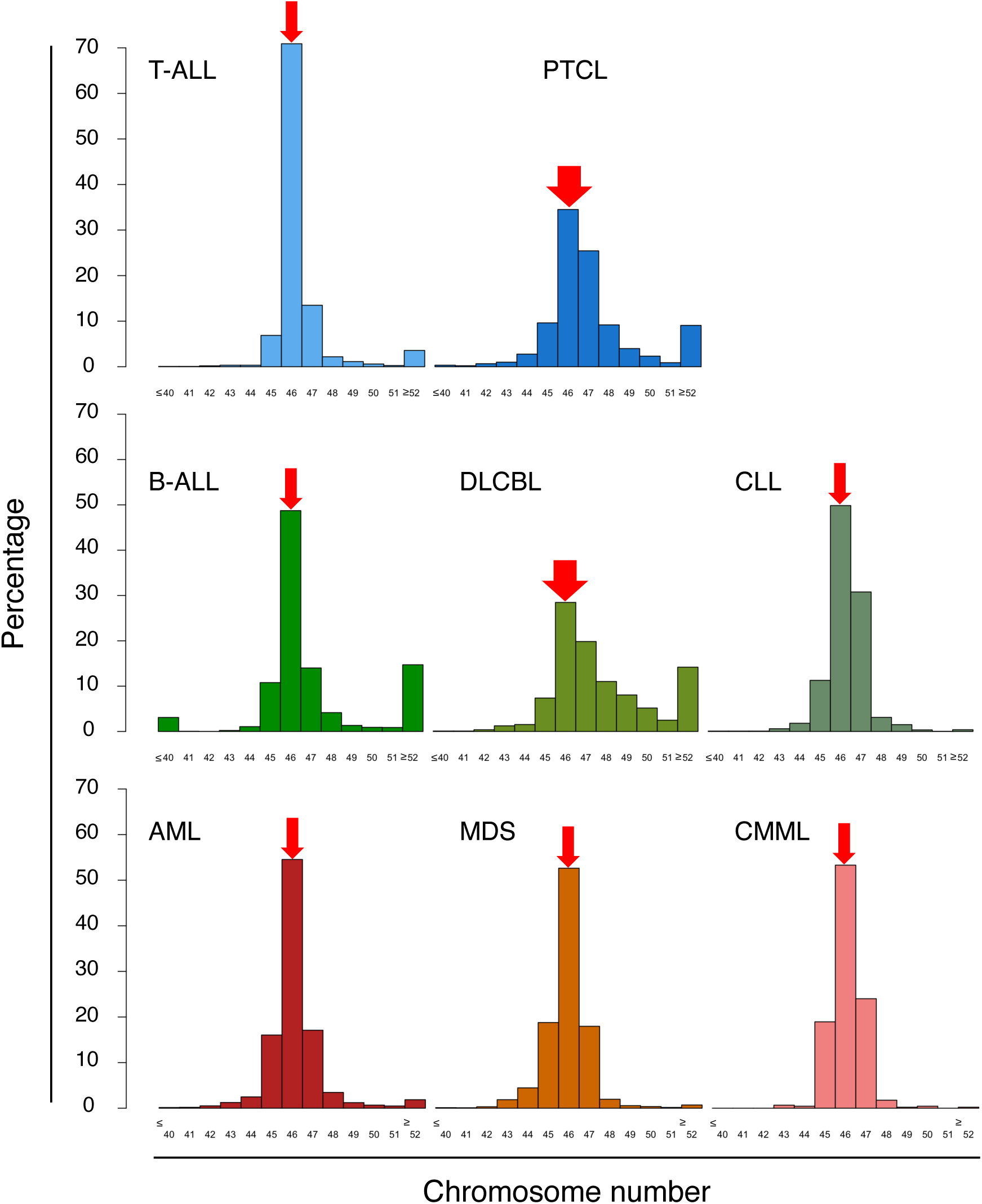
Chromosome number distributions across human hematologic malignancies. Percentage of cases with the indicated chromosome number, centered on the diploid chromosome number of 46 chromosomes, for the indicated hematologic malignancies. Histograms show chromosome number distributions for T-cell acute lymphoblastic leukemia (T-ALL), peripheral T-cell lymphoma (PTCL), B-cell acute lymphoblastic leukemia (B-ALL), diffuse large B-cell lymphoma (DLBCL), chronic lymphocytic leukemia (CLL), acute myeloid leukemia (AML), myelodysplastic syndrome (MDS), and chronic myelomonocytic leukemia (CMML). The red arrow at 46 chromosomes indicates the expected chromosome number for euploid cells. PTCL and DLBCL exhibited the lowest proportion of cases with 46 chromosomes, consistent with a greater prevalence of aneuploid karyotypes in these malignancies. Data were obtained from the Mitelman Database (*Mitelman Database of Chromosome Aberrations and Gene Fusions in Cancer (2026). Mitelman F, Johansson B, Mertens F and Paulsson K (Eds.),* https://mitelmandatabase.isb-cgc.org).

