## Supplementary Table S1 for "TET loss of function selects for aneuploidies that confer a proliferative advantage"

**Supplementary Table S1. Differentially expressed chromosome 17 genes in Tet2/3 DKO iNKT cells recovered after transplantation relative to WT iNKT cells.**

| Gene_name | log2FoldChange | Adjusted_pvalue | baseMean |
|---|---|---|---|
| Zfp57 | 2.815539007 | 0.025827474 | 14.3860367 |
| Gm33257 | 3.090606478 | 4.6618E-06 | 17.77451102 |
| Gnl1 | 0.669551159 | 6.83152E-09 | 2143.461334 |
| Prr3 | 0.591840205 | 8.239E-09 | 514.6921616 |
| Abcf1 | 0.746498035 | 6.37205E-07 | 3574.647584 |
| Mrps18b | 1.390547059 | 4.03654E-16 | 222.3109737 |
| 2310061I04Rik | 0.670350005 | 1.85498E-08 | 425.4050169 |
| Nrm | 1.540777505 | 1.9708E-18 | 546.8728109 |
| Tubb5 | 1.85870821 | 9.51853E-15 | 18513.59275 |
| Ddr1 | 2.11053068 | 0.000739113 | 1085.802454 |
| Gtf2h4 | 1.197148543 | 1.49555E-08 | 402.2769004 |
| Vars2 | 0.911776423 | 3.55529E-17 | 964.0617133 |
| Cchcr1 | 0.857158732 | 0.029162202 | 375.074312 |
| Atp6v1g2 | 1.11170029 | 0.003798536 | 80.08480805 |
| Prrc2a | 0.573453903 | 2.36736E-08 | 12445.79662 |
| Bag6 | 0.607920539 | 8.68811E-15 | 5321.627297 |
| Csnk2b | 0.581970761 | 0.000511381 | 681.0638752 |
| Clic1 | 0.56783341 | 1.15767E-10 | 6028.497324 |
| Msh5 | 1.660776518 | 4.98647E-05 | 30.48853693 |
| Lsm2 | 1.584269154 | 2.94838E-12 | 473.7520427 |
| Hspa1l | 1.856203739 | 3.08377E-07 | 41.94191957 |
| LOC115488834 | 1.982475948 | 0.039169661 | 6.283096042 |
| Snord52 | 2.166272331 | 0.008740873 | 6.933634603 |
| 1110038B12Rik | 2.458305564 | 2.2186E-109 | 645.6273303 |
| Ehmt2 | 0.510409687 | 0.02322635 | 4707.142116 |
| Zbtb12 | 1.983814602 | 7.81817E-19 | 172.1588549 |
| Cfb | 1.406506606 | 0.006699548 | 24.77374783 |
| Nelfe | 1.607868132 | 4.32404E-13 | 579.6382166 |
| Stk19 | 0.742709487 | 6.78286E-06 | 226.5557091 |
| C4a | 1.336163174 | 0.041423523 | 19.39781422 |
| BC051537 | 1.705740326 | 0.001438144 | 13.10670546 |
| Col11a2 | 1.090887192 | 0.000305797 | 284.1508028 |
| Rps18 | 0.912045036 | 4.98614E-14 | 1965.836102 |
| Wdr46 | 1.641314134 | 9.82154E-32 | 1118.234346 |
| Pfdn6 | 1.342012774 | 1.42678E-12 | 612.8779734 |
| BC051226 | 1.112990447 | 2.38488E-05 | 57.33651452 |
| Kifc1 | 1.630676648 | 1.02092E-08 | 318.5650298 |
| Cd320 | 2.12736923 | 7.06985E-22 | 225.3527244 |
| Ndufa7 | 0.649491486 | 7.39865E-08 | 765.3823429 |
| Rps28 | 1.07925414 | 8.83565E-17 | 671.4444318 |
| Kank3 | 1.470119196 | 4.59011E-06 | 55.50617809 |
| Angptl4 | 1.820831918 | 9.46947E-16 | 328.6424903 |
| Gm17251 | 0.835234122 | 0.022991789 | 38.1019598 |

| | | | |
|---|---|---|---|
| Srsf3 | 0.563556323 | 0.000147229 | 2221.725365 |
| Mapk13 | 5.186670252 | 0.024470249 | 2.378718893 |
| Mapk14 | 1.072715362 | 4.84181E-08 | 3669.856965 |
| Srpk1 | 0.927384557 | 1.26177E-37 | 4926.98861 |
| Fkbp5 | 1.979205141 | 7.60317E-09 | 3039.018765 |
| Tead3 | 2.749220374 | 0.008427617 | 6.752566986 |
| Rpl10a | 1.77002951 | 7.78107E-20 | 463.0179056 |
| Zfp523 | 0.780569365 | 1.42619E-10 | 664.931041 |
| Scube3 | 3.713093545 | 0.032641684 | 2.396393134 |
| Uhrf1bp1 | 1.043816602 | 2.77952E-09 | 389.1798254 |
| Snrpc | 1.097585247 | 1.80249E-07 | 245.3462709 |
| Pacsin1 | 0.980220348 | 0.001910562 | 1763.458137 |
| Rps10 | 0.767435372 | 5.34986E-07 | 2393.646374 |
| Nudt3 | 0.748573282 | 1.48217E-12 | 1894.46611 |
| Hmga1 | 2.071615488 | 1.03062E-19 | 531.3154962 |
| Tcf19 | 2.408933886 | 4.29876E-34 | 793.9224354 |
| AA388235 | 1.515008347 | 1.15623E-14 | 195.3446468 |
| Pot1b | 0.673532535 | 0.000599581 | 467.7528097 |
| Treml2 | 2.857327389 | 3.74242E-28 | 2051.660819 |
| Foxp4 | 1.095111247 | 6.42388E-09 | 1660.733266 |
| Mrps10 | 1.442712741 | 3.91781E-22 | 241.1149208 |
| Gm1976 | 2.494280657 | 1.08953E-20 | 80.9106883 |
| 2700099C18Rik | 3.325073554 | 1.42115E-17 | 84.11194091 |
| Msh6 | 1.645110562 | 2.07003E-13 | 1418.551322 |
| Msh2 | 1.475654112 | 2.94599E-34 | 2272.594419 |
| Epcam | 2.44700967 | 1.51924E-06 | 44.12117617 |
| Calm2 | 0.620984329 | 0.004802848 | 2080.226431 |
| Pigf | 0.666165199 | 0.0050717 | 89.76418682 |
| Epas1 | 2.488977318 | 0.004523254 | 1163.993204 |
| Srbd1 | 0.521499345 | 0.000351038 | 680.5835196 |
| Lrpprc | 1.193249533 | 2.529E-15 | 1857.967485 |
| Thada | 0.558871294 | 0.00383866 | 1196.416025 |
| Gm41640 | 0.937534471 | 0.0050717 | 72.66378773 |
| Haao | 1.414601137 | 0.003564319 | 25.81032189 |
| Gm46594 | 1.363588565 | 0.032652564 | 50.97924981 |
| Eml4 | 1.277569994 | 1.25512E-25 | 2793.050038 |
| Thumpd2 | 0.786246333 | 2.27249E-07 | 243.2348602 |
| Morn2 | 2.067709746 | 0.000102939 | 18.40532695 |
| Gemin6 | 2.439882585 | 8.88733E-13 | 107.148377 |
| Hnrnpll | 0.739597904 | 8.27426E-07 | 1558.601565 |
| Atl2 | 0.77807146 | 0.00256679 | 708.7577992 |
| Qpct | 1.313623138 | 0.000582501 | 1103.454038 |
| Prkd3 | 1.468893424 | 5.22045E-13 | 5690.793093 |
| Ndufaf7 | 0.980208504 | 8.25483E-07 | 372.9621907 |
| Cebpz | 0.943210458 | 3.92818E-19 | 1981.225595 |
| Cebpzos | 0.641888443 | 0.01128047 | 96.52652668 |
| Strn | 0.607046231 | 2.17256E-11 | 1420.793872 |
| Fam98a | 1.146385898 | 5.25975E-09 | 823.1289069 |

| | | | |
|---|---|---|---|
| Ttc27 | 1.515895141 | 2.85435E-36 | 786.6752532 |
| Yipf4 | 0.50788929 | 1.85222E-07 | 838.7954135 |
| Slc30a6 | 0.644122048 | 4.77291E-06 | 264.309373 |
| Spast | 0.562727199 | 1.22502E-05 | 1133.065192 |
| Dpy30 | 1.275204699 | 8.14884E-14 | 567.3943409 |
| Memo1 | 1.727755607 | 2.02353E-44 | 918.1527969 |
| Lclat1 | 1.772195328 | 9.25366E-32 | 751.4133631 |
| Togaram2 | 2.788725646 | 0.048166169 | 6.602005343 |
| Snord53 | 1.966236784 | 0.00431787 | 8.177611745 |
| Wdr43 | 1.292670557 | 1.36282E-25 | 2818.561139 |
| Trmt61b | 1.424961565 | 4.80359E-10 | 70.67915687 |
| Ndc80 | 2.729313622 | 9.90029E-19 | 473.3781629 |
| Smchd1 | 0.703751964 | 8.66297E-10 | 3687.018372 |
| Lpin2 | 0.841444975 | 0.000484373 | 1053.531558 |
| Myom1 | 1.431408973 | 0.013063973 | 15.66894288 |
| Tgif1 | 2.141109146 | 5.65127E-11 | 504.1104551 |
| C030034I22Rik | 0.952149207 | 0.001617891 | 61.02403423 |
| Arhgap28 | 3.297162952 | 0.041019371 | 17.65751511 |
| Ddx11 | 1.208268335 | 4.34578E-16 | 880.9752889 |
| Washc1 | 0.578516105 | 1.71008E-06 | 1176.583149 |
| Ndufv2 | 1.141744258 | 1.12598E-13 | 1303.639875 |
| Rab31 | 1.780637748 | 0.002182948 | 163.1780685 |
| Txndc2 | 4.523366621 | 0.0463556 | 4.28132308 |
| Vapa | 0.754649409 | 2.03871E-12 | 1714.824936 |
| Trip10 | 1.226262694 | 1.47501E-05 | 157.7183179 |
| Gm11110 | 0.940772369 | 0.000477461 | 79.39343361 |
| Tubb4a | 1.208564987 | 3.544E-05 | 64.31141742 |
| Khsrp | 1.068762223 | 1.92901E-11 | 4002.15669 |
| Gtf2f1 | 0.778287426 | 3.47599E-06 | 1374.200291 |
| Clpp | 0.966222595 | 1.31081E-07 | 393.0671378 |
| Mllt1 | 1.071549868 | 5.2599E-17 | 1650.686059 |
| Acsbg3 | 1.113958338 | 0.00155714 | 32.9316857 |
| Rfx2 | 0.913625492 | 0.004128197 | 83.74274851 |
| Prr22 | 1.741034968 | 4.32389E-08 | 35.39387933 |
| Dus3l | 0.583373997 | 3.11952E-06 | 961.4318907 |
| Ndufa11 | 0.737779666 | 3.00375E-05 | 510.385425 |
| Lonp1 | 0.86929092 | 1.76685E-10 | 1475.589101 |
| Rpl36 | 0.73565157 | 0.002077405 | 93.72952343 |
| Micos13 | 0.68564437 | 8.58105E-08 | 744.7227844 |
| Gm32822 | 4.74785377 | 0.023406043 | 1.570776032 |
| Ptprs | 3.091873773 | 1.52955E-38 | 1912.394026 |
| Uhrf1 | 3.806437342 | 5.42582E-25 | 2170.154513 |
| Plin3 | 1.559797564 | 0.012091222 | 787.1015723 |
| Mydgf | 1.235822602 | 1.50711E-07 | 367.0683927 |
| Chaf1a | 3.435144654 | 7.00844E-20 | 786.0322033 |
| Mpnd | 0.659799577 | 0.000121034 | 806.3396037 |
| Stap2 | 1.840764068 | 0.040345892 | 9.468753903 |
| Sgo1 | 3.906656016 | 6.27579E-28 | 338.1907582 |

| | | | |
|---|---|---|---|
| Gm19585 | 0.570709051 | 0.017935481 | 1420.85611 |
| Tbc1d5 | 0.802844227 | 5.36569E-14 | 1101.940001 |
| Rftn1 | 0.587187374 | 2.17086E-05 | 3125.153124 |
| Mdfi | 3.593647626 | 0.00063266 | 8.664856635 |
| Tomm6os | 2.616990992 | 0.033218209 | 2.881728373 |
| Tomm6 | 1.044487515 | 5.61522E-09 | 1037.075563 |
| Usp49 | 1.3661185 | 0.000138082 | 34.88820039 |
| Med20 | 0.653893177 | 2.93417E-06 | 472.5829664 |
| Bysl | 0.818258063 | 5.22005E-05 | 644.4643916 |
| Rpl7l1 | 1.036141923 | 7.51386E-26 | 1621.835253 |
| 2310039H08Rik | 1.241883817 | 2.8167E-13 | 146.205307 |
| Pex6 | 0.554066493 | 0.002259542 | 1628.940176 |
| Ppp2r5d | 0.671726442 | 4.83153E-06 | 1471.377659 |
| Klhdc3 | 1.248007741 | 4.30117E-28 | 1634.84031 |
| Cul7 | 1.000356475 | 0.008156867 | 445.0398409 |
| Mrpl2 | 1.159264875 | 2.78898E-17 | 477.032596 |
| Ptk7 | 2.71404942 | 0.001406223 | 115.27743 |
| Dnph1 | 4.040766097 | 6.65291E-07 | 29.69069216 |
| Zfp318 | 2.788987003 | 5.01534E-10 | 831.1161735 |
| Dlk2 | 2.704891587 | 0.035027252 | 4.3335076 |
| Lrrc73 | 2.109149612 | 0.004761437 | 12.71238373 |
| Polr1c | 0.930518128 | 2.2706E-19 | 891.2062764 |
| Xpo5 | 1.030226932 | 5.26877E-35 | 2103.000838 |
| Polh | 0.793951949 | 4.48064E-05 | 496.4948435 |
| Mrps18a | 0.980473187 | 1.67054E-12 | 454.7864523 |
| Vegfa | 5.988340544 | 0.000605123 | 24.7488444 |
| Gm35456 | 6.362807191 | 0.000124103 | 4.442042001 |
| Mrpl14 | 0.590155707 | 0.006457845 | 232.3651851 |
| Slc29a1 | 2.794959929 | 3.17053E-29 | 2604.022554 |
| Hsp90ab1 | 1.813962305 | 2.5082E-18 | 51669.49424 |
| Slc35b2 | 0.622657059 | 1.15432E-06 | 829.1920753 |
| Aars2 | 0.791614859 | 3.57492E-08 | 434.2744684 |
| B230354K17Rik | 1.616675189 | 9.14086E-13 | 354.4876158 |
| Cdc5l | 0.546764512 | 6.58632E-05 | 1350.55288 |
| Supt3 | 1.612251061 | 8.508E-21 | 192.4860184 |
| Runx2 | 1.484768896 | 0.001596663 | 286.0300503 |
| Adgrf1 | 4.492626702 | 0.025189406 | 1.898503448 |
| Tnfrsf21 | 2.764266454 | 6.00937E-15 | 287.7742188 |
| Cd2ap | 1.531810998 | 9.9798E-20 | 914.650202 |
| Mmut | 0.786312456 | 2.60566E-15 | 500.5876495 |
| Cenpq | 0.753721451 | 5.38163E-06 | 386.4828396 |
| 9130008F23Rik | 3.862261674 | 0.034759257 | 1.811453523 |
| Hnrnpm | 0.553311511 | 0.008640561 | 5781.186589 |
| Zfp414 | 1.023367804 | 2.44368E-08 | 442.3175046 |
| Zfp101 | 0.804970645 | 0.000199315 | 182.4967669 |
| Zfp955a | 0.642168322 | 4.7031E-05 | 210.8534585 |
| Wiz | 0.955202028 | 7.6168E-18 | 1402.967187 |
| Rrp1b | 1.471624499 | 2.62536E-11 | 1383.074166 |

| | | | |
|---|---|---|---|
| U2af1 | 0.899703124 | 5.677E-06 | 330.6554533 |
| Cbs | 4.733339194 | 0.006728303 | 2.691936105 |
| Wdr4 | 1.411867558 | 4.84191E-43 | 715.4777755 |
| Dnah8 | 3.708068801 | 2.03125E-38 | 1962.644009 |
| 1700097N02Rik | 4.532723625 | 7.10227E-26 | 194.3852566 |
| Glo1 | 1.408920953 | 4.78147E-19 | 496.3297254 |
| Btbd9 | 0.583544392 | 6.12956E-12 | 844.4026946 |
| Mtch1 | 1.312402711 | 1.39672E-19 | 1459.018836 |
| BC004004 | 0.581799924 | 1.23452E-11 | 972.2538536 |
| Ppil1 | 2.002673479 | 1.74623E-12 | 1105.36972 |
| Uqcc2 | 1.001177886 | 9.20058E-07 | 478.7525805 |
| Bak1 | 0.797971177 | 5.20503E-06 | 1738.041144 |
| Syngap1 | 2.164974766 | 1.11368E-07 | 36.70694047 |
| Kifc5b | 1.929909932 | 3.56383E-10 | 104.0509402 |
| Bnip1 | 0.583335912 | 9.66166E-05 | 314.604568 |
| Ergic1 | 0.991313176 | 7.43329E-16 | 1639.837395 |
| Luc7l | 0.760221055 | 1.04566E-10 | 2144.354052 |
| Mrpl28 | 1.18677363 | 4.48676E-12 | 663.6800394 |
| Nme4 | 3.169735844 | 3.00366E-08 | 157.5295757 |
| Pigq | 0.862072941 | 4.06829E-24 | 1506.230785 |
| Mettl26 | 1.075757526 | 4.43528E-11 | 498.0662464 |
| Mcrip2 | 2.416881438 | 4.24052E-13 | 74.5542435 |
| Wdr90 | 2.129435445 | 9.48203E-39 | 753.2911994 |
| Rhot2 | 0.909751678 | 2.33321E-17 | 1794.608347 |
| Rhbdl1 | 1.316707906 | 2.4126E-06 | 146.0726755 |
| Stub1 | 0.575226654 | 2.11842E-05 | 317.1072338 |
| Jmjd8 | 0.575336118 | 0.000167047 | 251.8331491 |
| Wdr24 | 0.611309503 | 6.79899E-08 | 747.208528 |
| Metrn | 1.380442073 | 0.000216374 | 544.8579378 |
| Ccdc78 | 1.450712838 | 0.003986896 | 16.88262059 |
| Haghl | 0.852355905 | 0.000886141 | 231.6794269 |
| Rpusd1 | 1.201525928 | 2.72148E-13 | 239.3945745 |
| Chtf18 | 2.356301904 | 1.40029E-13 | 355.1246737 |
| Lmf1 | 1.337046589 | 3.82289E-26 | 377.7913639 |
| Cerox1 | 5.807938535 | 1.02316E-13 | 39.02029598 |
| BC003965 | 1.120964662 | 1.35762E-21 | 476.3002708 |
| Telo2 | 1.226647154 | 2.97422E-34 | 788.586174 |
| Ift140 | 1.124170886 | 1.53051E-29 | 1021.303753 |
| Jpt2 | 2.231661663 | 3.50889E-27 | 747.6171566 |
| Mrps34 | 0.773708372 | 5.64154E-08 | 535.832456 |
| Nubp2 | 0.933700691 | 1.33138E-13 | 1020.493388 |
| Fahd1 | 1.78811451 | 3.6482E-07 | 54.98739175 |
| Hagh | 0.529393874 | 0.001647156 | 239.6656113 |
| Msrb1 | 0.517735897 | 0.001762282 | 308.7095238 |
| Ndufb10 | 0.646569103 | 0.000848959 | 1010.578839 |
| Rps2 | 1.173547885 | 1.01721E-11 | 3047.949656 |
| Tbl3 | 1.374216539 | 3.25369E-17 | 1019.217512 |
| Gfer | 1.009801452 | 6.84088E-11 | 272.4441329 |

| | | | |
|---|---|---|---|
| Slc9a3r2 | 3.90380247 | 4.95976E-06 | 13.1642183 |
| Nthl1 | 1.276367075 | 0.000717698 | 43.65834228 |
| Rab26os | 1.762046604 | 1.56963E-06 | 36.21817576 |
| Mlst8 | 1.005848887 | 7.45351E-23 | 427.7994095 |
| Pgp | 0.952613267 | 0.00050673 | 757.4810101 |
| Eci1 | 0.760318034 | 0.000213858 | 296.082586 |
| Rnps1 | 1.248827172 | 8.72743E-19 | 616.0353737 |
| D330041H03Rik | 1.15761971 | 0.003030842 | 36.65612556 |
| Abca3 | 1.473543435 | 3.94678E-08 | 855.6235959 |
| Abca17 | 4.876382347 | 0.00352923 | 3.48481609 |
| Ccnf | 3.413068686 | 4.83926E-17 | 555.1057408 |
| Tedc2 | 1.320274537 | 1.79033E-09 | 152.4231199 |
| Tbc1d24 | 2.648709486 | 1.50896E-28 | 119.5976693 |
| Kctd5 | 1.097417384 | 4.81774E-13 | 678.8732748 |
| Prss32 | 4.561001392 | 6.16083E-05 | 9.187687068 |
| Elob | 0.767098173 | 0.000152332 | 882.9678856 |
| Srrm2 | 0.56955562 | 0.020497524 | 21474.50243 |
| Flywch1 | 1.00051033 | 8.79987E-09 | 639.1403749 |
| Paqr4 | 2.699480237 | 3.70361E-13 | 141.9145608 |
| Pkmyt1 | 1.72658775 | 7.55638E-14 | 492.8926711 |
| Tnfrsf12a | 0.942817621 | 0.001703824 | 46.15295053 |
| Thoc6 | 1.62706848 | 7.85948E-28 | 420.5145962 |
| Zfp213 | 0.806405838 | 1.69794E-12 | 312.5028829 |
| 6330415G19Rik | 1.337999876 | 0.029320232 | 13.71474038 |
| Zfp40 | 0.959119814 | 2.00044E-05 | 199.9148817 |
| Gm7072 | 0.54375648 | 0.007097632 | 514.4132698 |
| Zfp947 | 1.421721749 | 0.033045276 | 17.65767533 |
| Zfp995 | 1.031524124 | 0.001325217 | 172.9226347 |
| Gm30531 | 2.624770897 | 0.033328053 | 3.238612138 |
| Zfp948 | 2.27136201 | 7.36723E-11 | 104.0001163 |
| Zfp52 | 1.562796539 | 7.29983E-16 | 546.6536153 |
| Ppp2r1a | 0.683035195 | 1.29589E-21 | 5345.810349 |
| Vmn2r100 | 8.313550647 | 1.41863E-05 | 28.28051864 |
| Vmn2r95 | 5.714872894 | 0.011283359 | 11.92456212 |
| Has1 | 3.033743017 | 0.00133953 | 13.17583968 |
| Spaca6 | 3.30090881 | 2.33096E-05 | 35.35977909 |
| Lnpep | 0.746767028 | 4.70499E-05 | 2849.447069 |
| Riok2 | 1.239450692 | 2.21205E-23 | 673.3871791 |
| Gm5165 | 0.618344591 | 0.033784103 | 83.09181008 |
| Chd1 | 0.664933544 | 4.15181E-12 | 3082.77727 |
| Psmb1 | 0.533071812 | 8.16074E-06 | 2734.627376 |
| Phf10 | 1.071406681 | 1.52086E-15 | 693.6635578 |
| Thbs2 | 7.270950652 | 0.001927265 | 11.64482193 |
| Smoc2 | 5.67865641 | 0.005912582 | 4.878495309 |
| Dact2 | 8.639181499 | 0.00155714 | 24.94561764 |
| Sod2 | 0.970744112 | 2.05848E-10 | 980.6341512 |
| Acat2 | 1.316850375 | 0.000504002 | 81.75662443 |
| Tcp1 | 1.512890956 | 6.14872E-13 | 4783.912747 |

| | | | |
|---|---|---|---|
| Mrpl18 | 1.070650864 | 7.35122E-06 | 888.2653121 |
| Slc22a3 | 1.148872266 | 0.046807642 | 521.5727755 |
| 4732491K20Rik | 2.743757174 | 0.00353002 | 7.276228219 |
| Agpat4 | 4.22576469 | 3.63224E-27 | 194.8831133 |
| Gm16702 | 4.282914978 | 3.72312E-21 | 112.6450791 |
| T2 | 4.538159598 | 2.15937E-07 | 17.1581484 |
| Gm8465 | 3.361385655 | 0.009464245 | 34.80453279 |
| Ccr6 | 3.918788739 | 0.005629205 | 305.8258805 |
| Cep43 | 1.77551089 | 1.11244E-40 | 701.1207617 |
| Gm41523 | 2.097274894 | 0.000605348 | 12.52478538 |
| Rsph3a | 1.092141223 | 0.015772279 | 27.54923752 |
| Tagap1 | 1.157103752 | 5.71289E-11 | 288.955887 |
| Ezr | 0.562809201 | 7.48661E-07 | 6607.693065 |
| Dynlt1f | 1.407097036 | 8.91681E-07 | 68.29896656 |
| Tmem181b-ps | 0.900759497 | 0.000241139 | 355.0730352 |
| Dynlt1a | 2.418602174 | 0.000607064 | 12.62732886 |
| Tulp4 | 0.5317807 | 0.013364824 | 574.2924901 |
| Gtf2h5 | 1.238361249 | 2.20371E-09 | 517.9555275 |
| Serac1 | 1.500896884 | 2.26793E-10 | 134.3896247 |
| Tmem242 | 1.217703479 | 8.1109E-29 | 441.3157107 |
| Tfb1m | 0.735659719 | 0.000270627 | 129.5946828 |
| Scaf8 | 0.537119967 | 4.78337E-18 | 2308.670869 |
| A230045G11Rik | 0.724941463 | 1.5676E-05 | 302.3691413 |
| Pisd-ps2 | 1.39117178 | 0.024732891 | 28.07361934 |
| Flot1 | -0.719784687 | 0.000694066 | 602.7909301 |
| Ier3 | -4.776801716 | 5.20413E-14 | 38.08470754 |
| Lta | -1.062727953 | 0.046891525 | 232.8378597 |
| Ltb | -1.126585453 | 1.92601E-05 | 5696.211715 |
| D17H6S53E | -0.674624042 | 0.000518494 | 294.0249916 |
| Ly6g5b | -2.246937378 | 2.85485E-07 | 553.7756274 |
| Neu1 | -0.879978847 | 0.040025378 | 1367.976307 |
| Prrt1 | -2.48947636 | 0.000925335 | 1267.348676 |
| Tap2 | -0.879265685 | 1.62511E-11 | 4733.252432 |
| Tap1 | -1.614922698 | 1.57087E-30 | 4768.386489 |
| Psmb9 | -1.241185734 | 7.05708E-14 | 1376.396904 |
| Tapbp | -0.79173601 | 2.98102E-06 | 7055.138487 |
| Smim40 | -5.512692739 | 0.000192927 | 8.964770161 |
| Bnip5 | -1.736475054 | 0.000581869 | 40.84287796 |
| Grm4 | -5.619236781 | 2.80224E-08 | 26.08092527 |
| Gm10505 | -2.041656755 | 3.1792E-08 | 83.69198718 |
| Unc5cl | -8.145069421 | 6.56913E-22 | 147.378303 |
| Apobec2 | -2.803023647 | 9.49541E-06 | 23.72175279 |
| Ttc7 | -0.798194907 | 4.53781E-08 | 2458.359511 |
| Socs5 | -0.534511091 | 6.48976E-05 | 527.7465235 |
| Rhoq | -1.612871581 | 7.0391E-27 | 249.4469756 |
| Prkce | -2.135034698 | 0.000151554 | 250.8329942 |
| Dync2li1 | -2.338070196 | 1.96362E-05 | 21.27260351 |
| Zfp36l2 | -1.597173361 | 7.65838E-05 | 9144.992061 |

| | | | |
|---|---|---|---|
| Galm | -4.696792681 | 8.78313E-29 | 171.3275311 |
| 4933407O12Rik | -0.674182317 | 0.041858714 | 55.61874075 |
| Rmdn2 | -1.49775295 | 2.19609E-06 | 352.7264075 |
| Gm17315 | -2.470540582 | 0.013478295 | 5.720404509 |
| Cdc42ep3 | -0.860179508 | 0.000277135 | 979.4668193 |
| Crim1 | -1.266237485 | 6.65995E-09 | 788.3059552 |
| Ltbp1 | -4.521945302 | 0.034420312 | 1.537105292 |
| Xdh | -2.742294742 | 1.91599E-06 | 667.9728856 |
| Ehd3 | -1.383092197 | 0.000579524 | 2219.942778 |
| Ypel5 | -0.564231572 | 0.011960619 | 817.7273362 |
| Ankrd12 | -0.650177274 | 0.000320621 | 997.1507206 |
| Nudt12 | -4.007606858 | 0.000145633 | 10.56623583 |
| Tnfsf14 | -0.976848553 | 0.000427021 | 1676.513653 |
| Satb1 | -1.279061157 | 1.27398E-12 | 9223.589988 |
| Ccnd3 | -1.46698419 | 6.14089E-10 | 2302.550021 |
| AI661453 | -6.906258485 | 1.28083E-07 | 25.88499268 |
| Bicral | -0.765887288 | 3.07997E-05 | 1302.428531 |
| Cul9 | -4.880229925 | 3.68584E-24 | 392.6117989 |
| Gtpbp2 | -0.634620285 | 0.000493866 | 2417.919527 |
| Nfkbie | -1.739438693 | 1.94689E-17 | 621.9865438 |
| Tmem151b | -2.469935313 | 0.026726862 | 16.29821168 |
| Clic5 | -2.522556823 | 0.023991884 | 13.05145865 |
| Gm8909 | -3.245671556 | 0.000321071 | 18.80583908 |
| Cyp4f17 | -6.18811016 | 5.98699E-17 | 45.87635557 |
| Pglyrp2 | -0.824842769 | 9.61127E-18 | 2750.878537 |
| Gm41569 | -1.068295344 | 7.54279E-06 | 115.9149351 |
| Rasal3 | -0.673397686 | 5.17507E-09 | 8248.467954 |
| Slc37a1 | -1.52089461 | 2.50796E-21 | 546.6960117 |
| Gm30139 | -1.03259778 | 0.01728982 | 20.96917935 |
| Rsph1 | -4.233912254 | 0.000295365 | 10.09612471 |
| Ubash3a | -0.974864518 | 0.04508075 | 550.0893866 |
| Abcg1 | -1.866918283 | 1.12385E-44 | 1670.284563 |
| Tbc1d22b | -0.538913607 | 1.84312E-08 | 1970.30563 |
| Pim1 | -1.210556597 | 5.14525E-05 | 1277.254605 |
| Gm26885 | -3.245647682 | 0.00439352 | 8.184156042 |
| Lemd2 | -0.504874615 | 0.000120773 | 860.1812467 |
| Phf1 | -2.418330975 | 5.72615E-13 | 1527.359351 |
| Crebrf | -0.961973188 | 0.003182242 | 1004.361529 |
| Dusp1 | -3.740548043 | 1.77655E-05 | 4719.017577 |
| Gm34455 | -3.65340796 | 0.016933686 | 4.539657951 |
| Rgs11 | -1.186415863 | 0.049535424 | 85.18302442 |
| Pgap6 | -0.598561568 | 1.661E-06 | 578.6764036 |
| Baiap3 | -3.600271767 | 1.33245E-10 | 337.0679639 |
| Dnase1l2 | -1.276933566 | 8.73915E-05 | 74.01456895 |
| Prss30 | -3.961242096 | 0.00569742 | 4.09153001 |
| Zfp53 | -0.519529812 | 0.000556352 | 474.0411437 |
| Gm52290 | -6.60745236 | 9.54615E-07 | 16.13392422 |
| Vmn2r96 | -2.443104444 | 0.033073829 | 5.496002628 |

| | | | |
|---|---|---|---|
| Afdn | -3.807516213 | 5.16773E-07 | 27.31130092 |
| C030013G03Rik | -2.640054261 | 0.036994667 | 3.469251501 |
| Rps6ka2 | -1.888784134 | 0.000226396 | 57.12439261 |
| Gm2808 | -2.264506615 | 0.017080214 | 10.25115279 |
| Gm46591 | -2.01124853 | 0.033308774 | 8.572545517 |
| H2-M3 | -1.066569804 | 7.93853E-26 | 576.7639663 |
| H2-M5 | 3.177724378 | 0.008823152 | 18.420406 |
| H2-T22 | -1.689510755 | 3.57589E-52 | 1349.611822 |
| H2-T23 | -1.413199605 | 4.37662E-14 | 2193.145702 |
| H2-T24 | -1.562153359 | 3.29897E-10 | 745.0281381 |
| H2-Q7 | -4.003239661 | 2.0696E-61 | 2192.87565 |
| H2-Q6 | -4.53998306 | 2.82481E-11 | 3333.615516 |
| H2-Q4 | -3.001406636 | 5.28819E-31 | 3349.809287 |
| H2-D1 | -1.948322137 | 1.30136E-34 | 18833.9401 |
| H2-Eb2 | 5.948586563 | 1.05653E-19 | 368.3678171 |
| H2-Ab1 | 2.729218306 | 0.044171677 | 105.1225387 |
| H2-DMa | -0.786558309 | 0.004022176 | 345.704121 |
| H2-Oa | 2.491147291 | 8.21024E-58 | 377.029328 |
| H2-K1 | -2.871562106 | 1.78066E-73 | 26219.01097 |
| H2-K2 | -1.06267144 | 0.000952165 | 363.9698524 |
| H2-Q5 | -4.397892478 | 4.76256E-32 | 305.8615017 |
| H2-T10 | -1.552088685 | 5.35213E-18 | 361.0868264 |
| H2-T-ps | 0.946886748 | 3.36269E-11 | 292.7451125 |
