## Supplementary Table S2 for "TET loss of function selects for aneuploidies that confer a proliferative advantage"

**Supplementary Table S2. Gene ontology (GO) categories and associated genes enriched among upregulated chromosome 17 genes in transplanted Tet2/3 DKO iNKT cells.**

| GO ID | Description | Upregulated genes within GO term |
|---|---|---|
| GO:0008380 | RNA splicing | Lsm2, Srsf3, Srpk1, Snrpc, Gemin6, Hnrnpll, Khsrp, Usp49, Cdc5l, Supt3, Hnrnpm, Rrp1b, U2af1, Ppil1, Luc7l, Rnps1, Srrm2, Thoc6 |
| GO:0034470 | ncRNA processing | Wdr46, Rps28, Srsf3, Rpl10a, Thada, Thumpd2, Wdr43, Dus3l, Bysl, Rpl7l1, Rrp1b, Wdr4, Rpusd1, Tbl3, Riok2, Gtf2h5, Tfb1m |
| GO:0007059 | Chromosome segregation | Tubb5, Bag6, Msh5, Ehmt2, Kifc1, Srpk1, Eml4, Ndc80, Washc1, Uhrf1, Sgo1, Cenpq, Kifc5b, Chtf18, Ppp2r1a, Riok2, Phf10 |
| GO:0022613 | Ribonucleoprotein complex biogenesis | Lsm2, Wdr46, Rps28, Srpk1, Rpl10a, Snrpc, Gemin6, Wdr43, Bysl, Rpl7l1, Rrp1b, Luc7l, Rpusd1, Tbl3, Riok2, Gtf2h5, Tfb1m |
| GO:0098813 | Nuclear chromosome segregation | Bag6, Msh5, Ehmt2, Kifc1, Eml4, Ndc80, Washc1, Uhrf1, Sgo1, Cenpq, Kifc5b, Ppp2r1a, Riok2, Phf10 |
| GO:0140053 | Mitochondrial gene expression | Mrps18b, Ndufa7, Mrps10, Lrpprc, Mrpl2, Mrps18a, Mrpl14, Aars2, Uqcc2, Mrpl28, Mrps34, Mrpl18, Tfb1m |
| GO:0032543 | Mitochondrial translation | Mrps18b, Ndufa7, Mrps10, Lrpprc, Mrpl2, Mrps18a, Mrpl14, Aars2, Uqcc2, Mrpl28, Mrps34, Mrpl18 |
| GO:0006364 | rRNA processing | Wdr46, Rps28, Rpl10a, Wdr43, Bysl, Rpl7l1, Rrp1b, Rpusd1, Tbl3, Riok2, Gtf2h5, Tfb1m |
| GO:0030490 | Maturation of SSU-rRNA | Wdr46, Rps28, Wdr43, Bysl, Tbl3, Riok2, Gtf2h5 |
