## Supplementary Table S3 for "TET loss of function selects for aneuploidies that confer a proliferative advantage"

**Supplementary Table S3. Expression of major histocompatibility complex (MHC) genes located on chromosome 17 in Tet2/3 DKO iNKT cells relative to WT iNKT cells.**

| Gene_names | log2FoldChange | Adjusted_pvalue | baseMean |
|---|---|---|---|
| H2-M2 | 1.671114793 | 0.569449399 | 1.795043054 |
| H2-M10.2 | 3.646594997 | 0.15108804 | 1.448137973 |
| H2-T3 | -3.378993576 | 0.133766886 | 1.762655326 |
| H2-Bl | -2.525781999 | 0.187579156 | 2.343364593 |
| H2-Q2 | -0.379515835 | 0.778732237 | 52.96301984 |
| H2-Q1 | -1.329693403 | 0.361892861 | 4.262527022 |
| H2-Aa | 1.710405002 | 0.182660585 | 73.95729732 |
| H2-Ob | -0.960440029 | 0.224680386 | 46.97779167 |
| H2-DMb1 | 1.347853617 | 0.257075941 | 9.312409074 |
| H2-DMb2 | 0.887455582 | 0.454772052 | 13.12085398 |
| H2-DMa | -0.786558309 | 0.004022176 | 345.704121 |
| H2-Ke6 | -0.06113411 | 0.826918343 | 942.1709714 |
| H2-T-ps | 0.946886748 | 3.36269E-11 | 292.7451125 |
| H2-M5 | 3.177724378 | 0.008823152 | 18.420406 |
| H2-Ab1 | 2.729218306 | 0.044171677 | 105.1225387 |
| H2-Oa | 2.491147291 | 8.21024E-58 | 377.029328 |
| H2-M3 | -1.066569804 | 7.93853E-26 | 576.7639663 |
| H2-T22 | -1.689510755 | 3.57589E-52 | 1349.611822 |
| H2-T23 | -1.413199605 | 4.37662E-14 | 2193.145702 |
| H2-T24 | -1.562153359 | 3.29897E-10 | 745.0281381 |
| H2-Q4 | -3.001406636 | 5.28819E-31 | 3349.809287 |
| H2-K2 | -1.06267144 | 0.000952165 | 363.9698524 |
| H2-Q5 | -4.397892478 | 4.76256E-32 | 305.8615017 |
| H2-T10 | -1.552088685 | 5.35213E-18 | 361.0868264 |
| H2-Q7 | -4.003239661 | 2.0696E-61 | 2192.87565 |
| H2-Q6 | -4.53998306 | 2.82481E-11 | 3333.615516 |
| H2-D1 | -1.948322137 | 1.30136E-34 | 18833.9401 |
| H2-Eb2 | 5.948586563 | 1.05653E-19 | 368.3678171 |
| H2-K1 | -2.871562106 | 1.78066E-73 | 26219.01097 |
